# CryoET Reveals Organelle Phenotypes in Huntington Disease Patient iPSC-Derived and Mouse Primary Neurons

**DOI:** 10.1101/2022.03.26.485912

**Authors:** Gong-Her Wu, Charlene Smith-Geater, Jesús G. Galaz-Montoya, Yingli Gu, Sanket R. Gupte, Ranen Aviner, Patrick G. Mitchell, Joy Hsu, Ricardo Miramontes, Keona Q. Wang, Nicolette R. Geller, Cristina Danita, Lydia-Marie Joubert, Michael F. Schmid, Serena Yeung, Judith Frydman, William Mobley, Chengbiao Wu, Leslie M. Thompson, Wah Chiu

## Abstract

Huntington’s Disease (HD) is caused by an expanded CAG repeat in the huntingtin gene, yielding a Huntingtin protein with an expanded polyglutamine tract. Patient-derived induced pluripotent stem cells (iPSCs) can help understand disease; however, defining pathological biomarkers is challenging. Here, we used cryogenic electron tomography to visualize neurites in HD patient iPSC-derived neurons with varying CAG repeats, and primary cortical neurons from BACHD, deltaN17-BACHD, and wild-type mice. In HD models, we discovered mitochondria with enlarged granules and distorted cristae, and thin sheet aggregates in double membrane-bound organelles. We used artificial intelligence to quantify mitochondrial granules, and proteomics to show differential protein content in HD mitochondria. Knockdown of Protein Inhibitor of Activated STAT1 ameliorated aberrant phenotypes in iPSC-neurons and reduced phenotypes in BACHD neurons. We show that integrated ultrastructural and proteomic approaches may uncover early HD phenotypes to accelerate diagnostics and the development of targeted therapeutics for HD.

## INTRODUCTION

Huntington’s disease (HD) is a progressive, fatal neurodegenerative disorder caused by a genetic mutation in the huntingtin gene (*HTT*)^1^. The mutation is an expansion of an N-terminal CAG repeat to 40 and above. This yields a mutated Huntingtin protein (mHTT) with an expanded polyglutamine (polyQ) tract that is pathogenic. Disease typically strikes in mid-life, lasting ∼10-15 years with ongoing progression of symptoms, which include cognitive decline, mood and personality disorders, and loss of motor control^2^. CAG length is correlated inversely with age of disease onset with repeats longer than ∼60 causing a juvenile form of HD^3, 4^. No disease-modifying treatments are available. Neuropathologically, degeneration of medium spiny neurons in the striatum and cortical atrophy serve as prominent manifestations^5^.

Methods to decipher the pathogenesis of neurodegenerative disorders are needed to identify biomarkers sensitive to clinical progression and to inform therapeutic trials. A potential source of such insights are patient-derived induced pluripotent stem cells (iPSCs)^6, 7^, which can be differentiated into multiple cell types, including neurons^8^ exhibiting disease phenotypes. Indeed, a number of HD-associated phenotypes have been recapitulated in neurons differentiated from HD iPSCs, including transcriptional dysregulation^4, 9–12^, bioenergetic deficits^13^, impaired neurodevelopment^9, 14, 15^, altered cell adhesion^10, 12^, impaired nucleocytoplasmic trafficking^16, 17^ and increased susceptibility to cell stressors^18^, among others^18^.

The propensity of mHTT to aggregate in neuronal cells is a hallmark of HD and leads to the appearance of large (micrometer scale) nuclear and neuritic inclusions^19^, as seen in mouse models^20^ and human^21^ post-mortem brains^19, 22^. Mutant HTT’s role in the potential impairment of autophagy may contribute to aberrant protein accumulation^23^. However, neither protein aggregation nor disruptions to protein homeostasis have been observed in iPSC-derived HD models unless treated with an inhibitor of the proteasome^24^, likely because these models represent early developmental stages where overt disease phenotypes are challenging to detect.

In parallel, other technological advances in cell, molecular and structural biology are poised to contribute to the goal of identifying early pathology biomarkers. For example, advances in cryogenic electron microscopy (cryoEM) and tomography (cryoET) have recently elucidated the structure of soluble HTT in complex with HAP40 at near atomic resolution^25^, the topology of mHTT-exon 1 and polyQ *in vitro* aggregates at nanometer resolution^26^, and have enabled visualization of the interactions between mHTT-exon 1 aggregates and other proteins and cellular compartments in transfected yeast^27^ and HeLa^28^ cells, and with molecular chaperones *in vitro*^29, 30^. Correlative light and traditional EM microscopy has also been used to image recruitment of mHTT-exon 1 to cytoplasmic aggregates within single membrane, vesicle-rich endolysosomal organelles^31^.

Herein, we used cryoET to visualize neurites from five human iPSC-derived neuronal cell lines endogenously expressing full-length mHTT with a range of CAG repeat lengths (Q18, Q53, Q66, Q77 and Q109). Using the same methods, we also studied mouse primary neurons from the BACHD model^32^ expressing full-length human mHTT, full-length human mHTT lacking the first 17 amino acids (deltaN17; dN17)^33^, and control WT mice. For all our samples, we examined subcellular organelles previously implicated in HD, namely mitochondria^34^ and autophagosomes^35^, and found marked changes in morphology as compared to controls. We then coupled these ultrastructural observations with mitochondrial proteomics and found changes in the levels of a number of mitochondrial proteins in HD samples. We also developed an artificial intelligence-based semi-automated 3D segmentation method to quantify changes in mitochondrial granule numbers and sizes.

Guided by combined ultrastructural and proteomic data, we explored the impact of genetic knockdown (KD) of a SUMO E3 ligase Protein Inhibitor of Activated STAT1 (PIAS1), a protein linked to maintenance of proteostasis and synaptic function in HD. In both systems, we observed rescue of mitochondrial morphology. Additionally, PIAS1 reduction abrogated aberrant aggregates in human HD iPSC-derived neuron autophagic organelles, findings consistent with prior studies showing that PIAS1 inhibition ameliorates HD pathology in mice and human iPSCs^36^.

Our investigations provide a platform with which to structurally evaluate *in situ* organelle phenotypes in thin regions of intact cells at nanometer resolution, in the presence and absence of potential therapeutics. The paradigm we propose emphasizes the ability to explore early disease manifestations and mechanisms in neurites of intact patient and mouse model-derived neurons, which serves as a proof of concept for the utility of cryoET as a structural readout to assess early HD phenotypes and to support preclinical evaluation of potential therapies.

## RESULTS

### Huntington’s disease patient-derived iPSCs differentiated into mature neurons on electron microscopy grids

Neurons differentiated from human iPSCs with pathological- and normal-length polyQ tracts in their *HTT* gene provide a platform to study different HD pathological states within their endogenous genetic context^4^. Here, we developed a robust protocol to differentiate iPSC-derived neurons with characteristics of medium spiny neurons on electron microscopy (EM) gold grids (**Supplementary Fig. 1**).

We first differentiated iPSCs to neural progenitors and then adapted our prior maturation protocol^4^ to allow the cells to grow directly on gold EM grids, eliminating the Matrigel matrix to minimize background densities, thereby maximizing contrast in cryoET images (**Supplementary Fig. 1a)**. The cells survived, differentiated and matured without Matrigel, producing axons and dendrites, and displaying normal neuronal morphology (**Supplementary Fig. 1b,c**). Differentiation of cells was also performed at half density (see Methods) to increase the likelihood of obtaining grids with only one cell per grid square, minimizing the potential for overlaps between cells while maximizing the number of areas suitable for cryoET imaging (**Supplementary Fig. 1c**). These modifications allowed iPSCs to differentiate into cells with medium spiny neuron-like characteristics, as validated by DARPP32 and CTIP2 co-staining^12^ (**Supplementary Fig. 1d)**. Cells were differentiated for 16 days, then plated on gold EM grids for terminal differentiation and maturation. After three more weeks (21 ± 2 days) of terminal differentiation, grids were vitrified by rapid plunging into liquid propane on day ∼37-39^37, 38^ to preserve the neurons on them in a near-native state without chemical fixative or metal stain.

### CryoET data showed mitochondria with abnormal cristae and enlarged granules in neurites of HD patient iPSC-derived neurons

The initial motivation to image intact human HD neurons using cryoET was to determine whether we could directly visualize at nanometer scale *in situ* aggregates of native mHTT (endogenous and untagged) that are not visible using other microscopy methods, as well as their surrounding subcellular components. A single iPSC-derived neuron is a micrometer-sized cell with a thick cell body and long, thin neurites. Since the electron beam cannot penetrate through the cell body of neurons, we extensively surveyed the structural features in the neurites of our HD cells by recording 2D cryoEM low-magnification projection images, from which we were able to identify potential regions of interest for subsequent higher magnification cryoET data collection (**Supplementary Fig. 2**).

Following an iterative search in many different areas on the cryoEM grids for all HD patient iPSC-derived (Q53, Q66, Q77 and Q109) neurons, we failed to detect large cytoplasmic inclusions or aggregates such as those formed by mHTT-exon 1 previously observed by cryoEM/cryoET *in vitro*^26, 29, 30, 39, 40^ and in transfected cells^28, 41^. However, this exhaustive examination did detect in numerous regions in the neurites of all samples abnormally large and dense, discrete, granular features as well as tangled aggregates, both within double membrane-bound compartments. Some of these compartments showed classic features of mitochondria (Fig. 1), as marked by an outer double membrane and the appearance of interior folded cristae, continuous with the inner membrane of the compartment (**Supplementary Video 1**).

The large and dense granular features inside mitochondria were consistently present in tens of tomograms of all HD cell lines (Q53, Q66, Q77 and Q109), and cristae were abnormal in most mitochondria in higher polyQ lines (Q66, Q77 and Q109)(**Fig. 1c-e**). In general, both phenotypes (namely, enlarged granules and disrupted cristae) became more pronounced with higher polyQ length. Importantly, these aberrant features were absent from mitochondria of control iPSC-derived neurons (Q18), though smaller features, consistent with normal mitochondrial granules, were readily observed (**Fig. 1a**). To facilitate 3D visualization, we used the convolutional neural network-based algorithm in EMAN2^42^ to annotate and segment these dense structures as well as other mitochondrial and subcellular features in surrounding areas, such as microtubules (**Fig. 1a-e, Supplementary video 1**). Of note, the enlarged granules in mitochondria of HD cells did not comprise homogeneous, smooth densities, but rather exhibited complex interwoven textures, occasionally displaying lower-density regions or voids (**Fig. 2**).

**Fig. 1.**
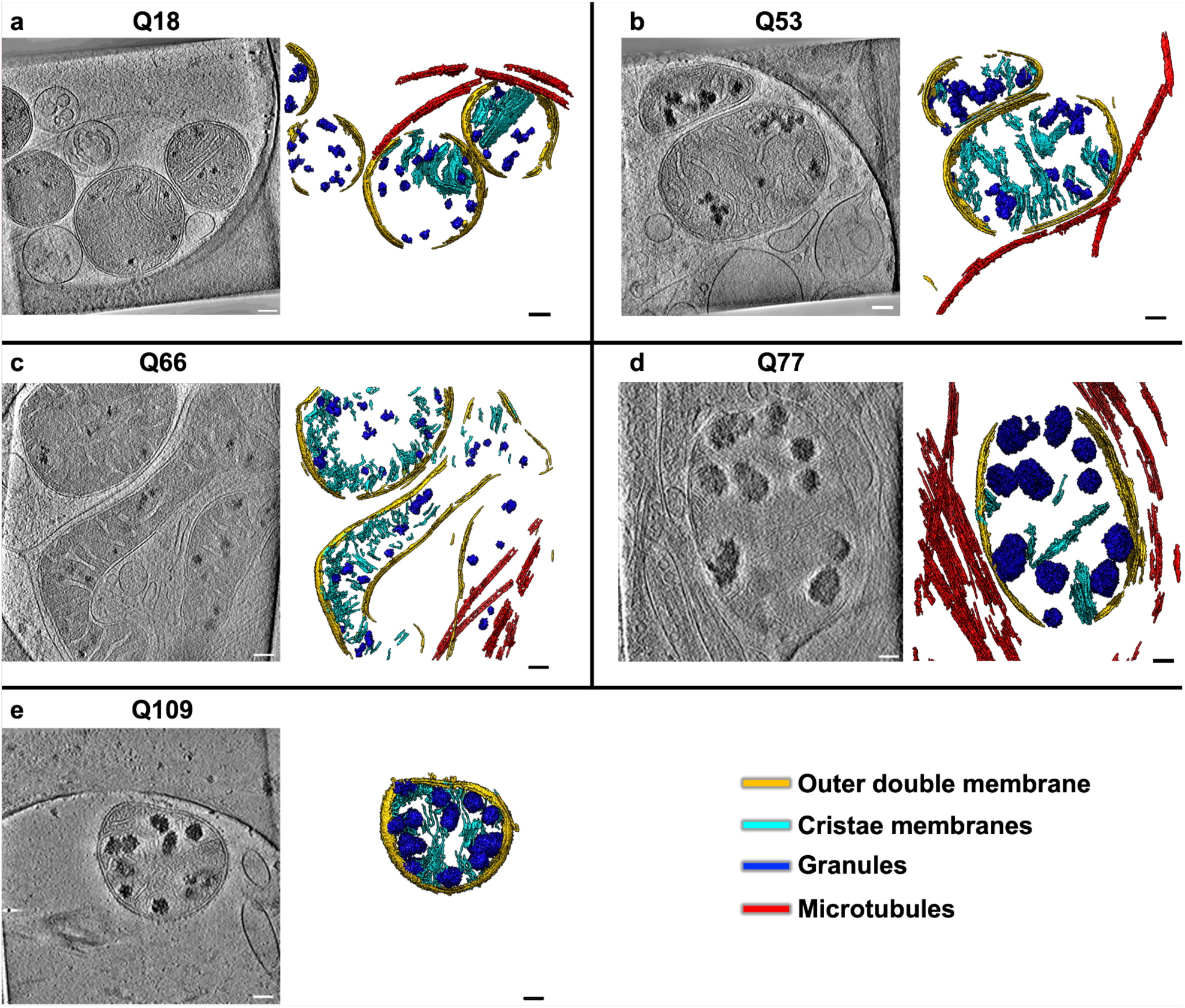
Mitochondria in neurites of HD patient iPSC-derived neurons exhibit altered morphology and contain enriched granules of varying size. Slices (∼1.4 nm thick) through selected regions of representative cryoET tomograms and corresponding segmentations of local features for a Q18, b Q53, c Q66, d Q77 and e Q109 human iPSC-derived neurons. Q53-Q109 reveal that mitochondria in HD patient-derived neurons have swollen cristae and contain enlarged granules compared to controls (Q18). Tomogram numbers: Q18=21, Q53=14, Q66=10, Q77=5 and Q109=37. Segmentation colors: red:microtubules, yellow:mitochondrial outer double-membranes, dark blue:granules, and cyan:cristae membranes. Scale bars = 100 nm.

**Fig. 2.**
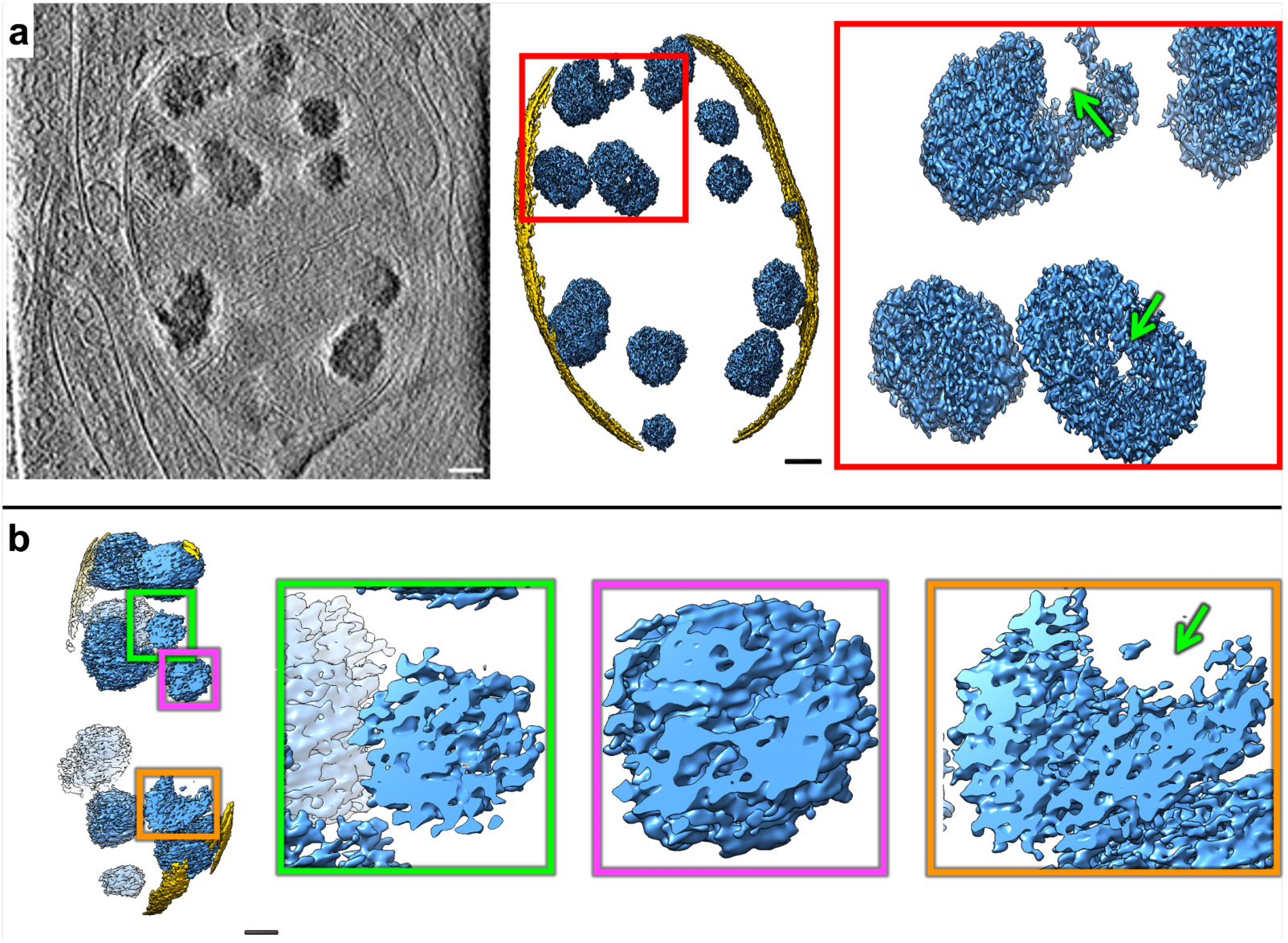
Mitochondria in HD neurites contain enlarged granules composed of tightly packed, heterogeneous densities. **a** Z**-**slice (∼1.4 nm thick) through a representative cryoET tomogram of a neurite in an HD patient iPSC-derived neuron (Q77) and corresponding segmentation of double membranes (yellow) and dense, granular densities inside (light blue). **b** Cutaway, oblique view of the segmentation in **a** and zoomed-in views of selected regions showing examples of interwoven densities in enlarged mitochondrial granules. Green arrows point to void regions. Scale bars = 100 nm. Segmentation colors: yellow:double membrane, light blue:mitochondrial granules.

### CryoET data showed mitochondria with abnormal cristae and enlarged granules in neurites of HD mouse model primary neurons

Next, we tested whether the abnormal ultrastructural features we observed in iPSC-neurons were also present in neurons derived from the well-studied BACHD mouse model^32^, which expresses full-length human mHTT with an expanded polyQ tract comprised of 97 mixed CAG-CAA repeats under the control of human regulatory sequences. We used cryoET to image neurites in primary cortical BACHD neurons and again found abnormally enlarged granules within mitochondria whose cristae were often disrupted compared to WT (**Fig. 3a,b**), similar to those seen in neurites of HD patient iPSC-derived neurons (**Fig. 1b-e**). Our data are evidence that neurons from both human and mouse HD models share the presence of enlarged granules and other changes in mitochondria, supporting the view that mHTT is responsible and raising the possibility that these morphological abnormalities could be used as diagnostic features. We also evaluated primary cortical neurons from BACHD transgenic mice expressing human mHTT lacking the first 17 N-terminal amino acids (dN17-BACHD)^33^. The N-terminus of HTT contains a putative mitochondrial membrane-targeting sequence that can form an amphipathic helix^43, 44^ characteristic of proteins transported into the mitochondria. This domain also promotes cytosolic localization of HTT^45^, potentially as a nuclear export sequence^46^ and is required for interaction with the subunit translocase of mitochondrial inner membrane 23 (Tim23), a mitochondrial protein import complex essential for protein import^47^. Phenotypes are more robust and progress more rapidly for dN17-BACHD than for BACHD mice^33^, with nuclear and endoplasmic reticulum (ER) mHTT localization for the former, as well as large nuclear inclusions visible by immunofluorescence, which are not observed in the BACHD model^33^. CryoET of neurites in dN17-BACHD neurons also showed severe distortions in most of their mitochondria (**Fig. 3c**). These observations therefore suggest that mHTT can disrupt mitochondria via polyglutamine repeat-dependent effects, such as those previously reported (using other techniques) to disrupt mitochondrial membranes^48^, independent of the N17 domain.

**Fig. 3.**
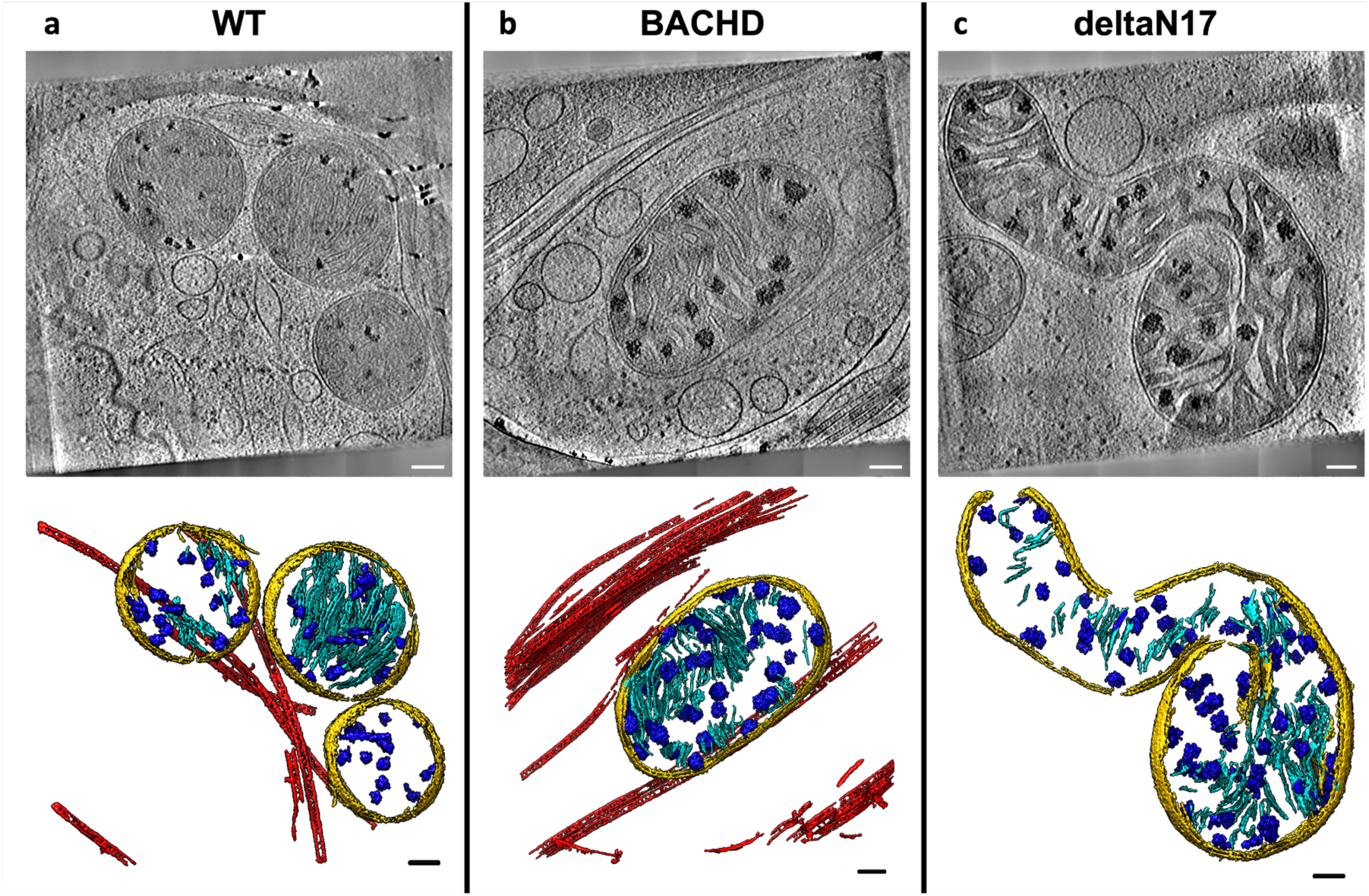
Mitochondria in neurites of HD mouse model neurons exhibit altered morphology and contain enlarged granules of varying size. Slices (∼1.4 nm thick) through selected regions of representative cryoET tomograms and corresponding segmentations of local features for **a** WT, **b** BACHD and **c** dN17 BACHD primary neurons reveal that neuronal mitochondria in HD mice have swollen cristae and contain enlarged granules compared to controls (WT). Tomogram number<u>s</u>: WT=31, BACHD=22 and deltaN17 BACHD=15. Segmentation colors: red:microtubules, yellow:mitochondrial double-membranes, dark blue:granules, and cyan:cristae. Scale bars = 100 nm.

### Neurites in HD patient iPSC-derived and mouse model neurons contain sheet-like aggregates in autophagic organelles

In addition to the dense, enlarged granules observed in mitochondria, we observed numerous, much larger aggregates in other membrane-bound compartments in both HD patient iPSC-derived and mouse primary neurons. While they appeared to be filamentous when viewed in two-dimensional (2D) z-slices through the tomograms, closer three-dimensional (3D) inspection, including the use of UCSF ChimeraX’s virtual reality (VR) graphics^49^, revealed that they are composed of densely interwoven slab-shaped, and sheet-like aggregates (**Fig. 4**, **Supplementary Video 2**). Indeed, these aggregates had a distinct sheet morphology, similar to that of intricate desert rose-like crystal of selenite, gypsum or barite found in the desert (hereafter called sheet aggregates) [https://en.wikipedia.org/wiki/Desert_rose_(crystal)].

**Fig. 4.**
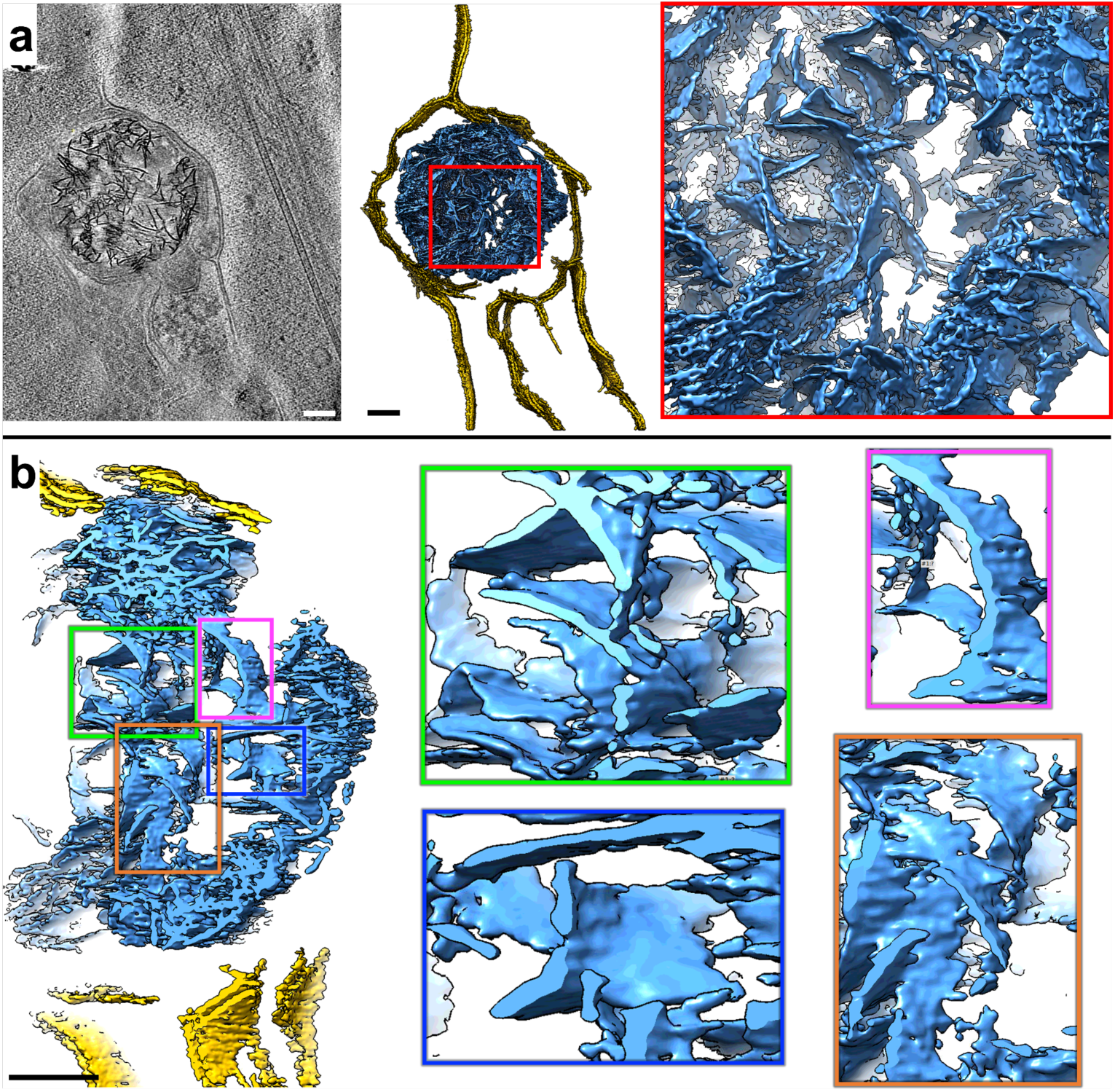
Neurites in HD cells contain double membrane-bound compartments with sheet aggregates composed of interwoven slabs and sheets. **a** Z-slice (∼1.4 nm thick) through a selected region showing a sheet aggregate in a representative cryoET tomogram of a neurite of a HD patient iPSC-derived neuron (Q66) and corresponding segmentation of double membranes (yellow) and aggregated densities inside (light blue). b Cutaway, oblique view of an enlarged region from the segmentation in a and further zoomed-in views of selected subregions showing examples of sheet-like areas within the aggregate. Scale bars = 100 nm. Segmentation colors: yellow:double membrane, light blue:sheet aggregate.

Importantly, these sheet aggregates were present in the neurites of all HD patient Qn neurons (**Fig. 5a**)(Q53, Q66, Q77 and Q109) as well as in those of HD mouse model neurons (**Fig. 5b**)(BACHD and deltaN17 BACHD). On the other hand, double-membrane bound compartments in neurites of control human iPSC-derived (Q18) (**Fig. 5c**) and mouse (WT) (**Fig. 5d**) neurons lacked these sheet aggregates.

**Fig 5.**
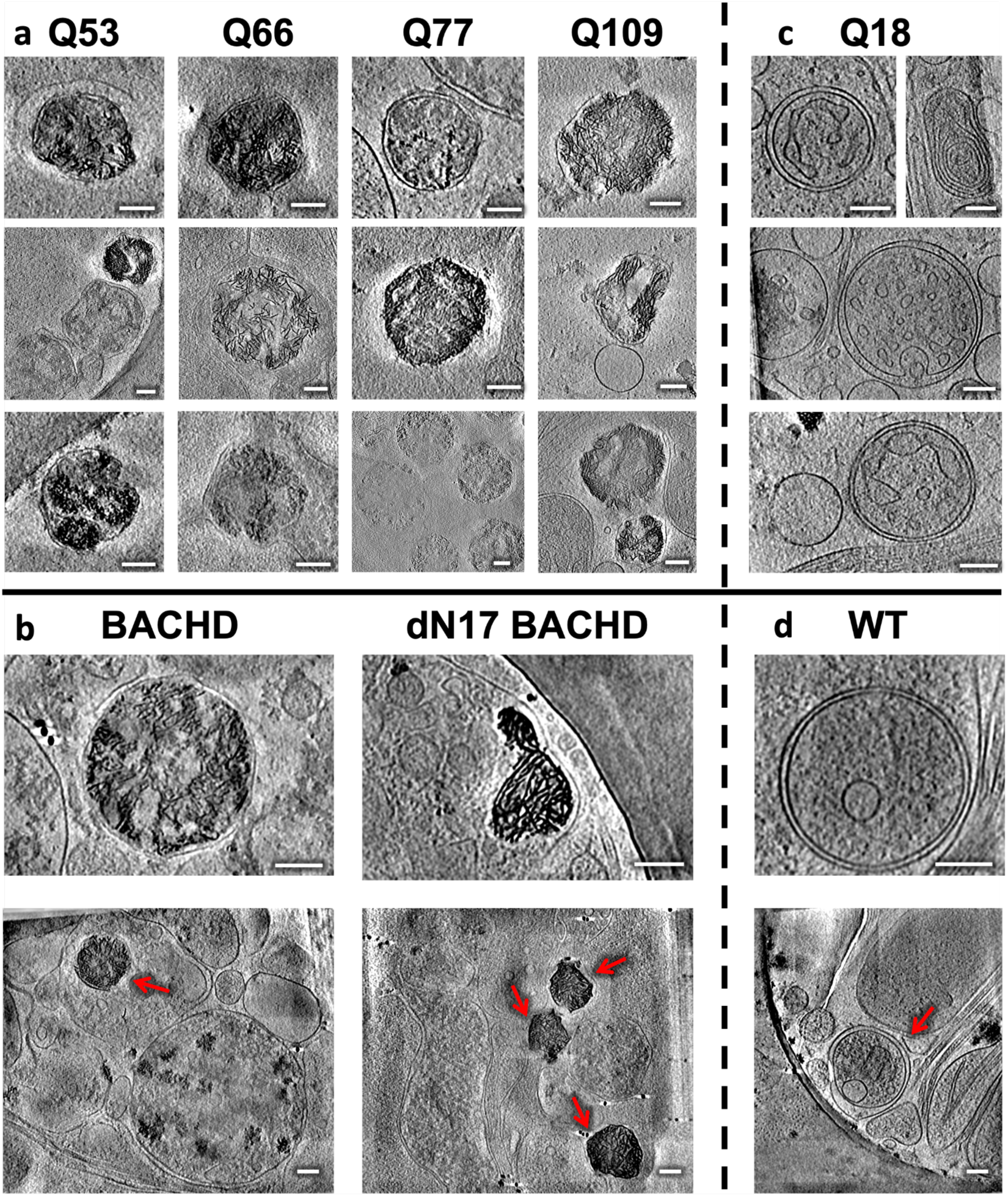
Neurites in HD patient iPSC-derived and mouse model primary neurons exhibit sheet aggregates within double membrane-bound organelles. Slices (∼1.4 nm thick) through selected regions of representative cryoET tomograms showing double membrane-bound compartments in neurites of HD **a** patient iPSC-derived (Q53, n=5; Q66, n=8; Q77, n=6; Q109, n=6) and **b** mouse model (BACHD, n=14; and dN17-BACHD, n=6) primary neurons, as well as normal **c** Q18 (n=4) and **d** WT (n=3) controls. Scale bars = 100 nm. The red arrows in full-frame views in **b** indicate double membrane-bound organelles.

Of note, the sheets in these aggregates were reminiscent of some regions in mHTT exon 1 and polyQ-only aggregates *in vitro*, shown to contain long, relatively flat or slightly curved ribbon-like^50^ or sheet-like^26^ morphologies as visualized by negative stain electron microscopy and cryoET, respectively. Interestingly, their thickness appeared relatively uniform, ∼2 nm, when visualized in our 3D tomograms with contrast transfer function correction and without downsampling or low-pass filtration. Importantly, we also collected higher-magnification cryoEM images of neurite regions with sheet aggregates and analyzed their Fourier transforms but did not detect any periodic arrangement in them.

Eukaryotic cells contain several characteristic double membrane-bound compartments including mitochondria, nucleus, and organelles in the autophagy pathway such as mitochondria-derived vesicles^51^, autophagosomes^52^, and amphisomes^52^. Here, the compartments containing the sheet aggregates (∼200 nm to ∼500 nm range in longest span) were much smaller than the nucleus (∼3-18 μm in diameter), and sometimes seemed to bind or merge with one another (**Supplementary Fig. 3a**). While they were most often similar in size to small mitochondria, and could possibly correspond to degenerating versions of this organelle or mitochondria-derived vesicles^51^, there is a possibility they may also correspond to other autophagic organelles with no visible cristae or to other molecular components targeted for autophagy^52^, suggesting alternative and possibly complementary or parallel biogenesis origins. For instance, autophagosomes participate in cellular pathways for degradation and clearance, and thus are possible candidates to contain these sheet aggregates. Supporting this interpretation, we found instances of sheet aggregates within double membrane-bound compartments fused with single membrane-bound compartments (**Supplementary Fig. 3b-d**), reminiscent of lysosomes, a picture strikingly similar to amphisomes, which result from fusion of autophagosomes with lysosomes^52^. On the other hand, in support of the degenerating mitochondria assignment, we observed features suggestive of nascent sheet aggregates in what appeared to be degenerating mitochondria (**Supplementary Fig. 3e**). To further characterize the nature of these features, we again used semi-automated, neural-networks-based annotation of the corresponding tomogram with EMAN2^42^, training on a few positive references (n=10) from a mature sheet aggregate only. Strikingly, the algorithm assigned the putative nascent sheet aggregate features in what appears to be a mitochondrion as belonging to the same type of feature as mature sheet aggregates, even though the networks were trained exclusively with mature sheet aggregate references (**Supplementary Fig. 3f**). This organelle could represent a degenerated mitochondrion bound with lysosomes, as these two organelles were recently discovered to interact directly via their membranes^53^. Indeed, this mitochondrion and a similar neighboring organelle are seen interacting with single membrane bound compartments (**Supplementary Fig. 3g**).

### Mitochondrial proteomics of human iPSC-derived neurons identified differentially expressed proteins, including those engaged in RNA binding

The accumulation of mitochondrial granules (**Fig. 2**), distinctly different from the sheet aggregates (**Fig. 4**), and the disruption of cristae observed in the neurites of HD neurons, are consistent with impaired mitochondrial function and bioenergetics previously described for HD^34, 54^. This phenotype could represent aberrant accumulation of RNA granules^55^, suggesting RNA processing and RNA quality control deficits, and/or aberrant protein quality control due to disrupted protein import arising from the presence of mHTT^47^. To investigate potential mechanisms underlying the abnormal enlargement of mitochondrial granules, we performed liquid chromatography tandem mass spectrometry-based proteomic analysis on mitochondria isolated from HD patient iPSC-derived neurons (Q109) and controls (Q18) since the former represents the most aggressive variant among our HD patient iPSC-derived samples (**Fig. 6**). Mitochondria were isolated by using magnetically labeled anti-TOM22 microbeads^56^. Isolated HD mitochondria looked similar in cryoEM projection images and reconstructed cryoET tomograms (**Fig. 6a**) to those in neurites containing enlarged granules (**Fig. 1e**). In quality control experiments, the mitochondrial isolation method was assessed via Western analysis on each cell fraction; this yielded a fraction with enrichment of mitochondria (using ATPB protein levels as a proxy) with some minor contamination from other organelles (LC3 levels were used as a proxy for autophagosomes, and CTIP2 levels as a proxy for nuclei) (**Supplementary Fig. 4a**). Consistent with successful isolation of mitochondria previously, our proteomic datasets were enriched for Gene Ontology (GO) terms related to mitochondrial functions (**Supplementary Fig. 4b**).

**Fig. 6.**
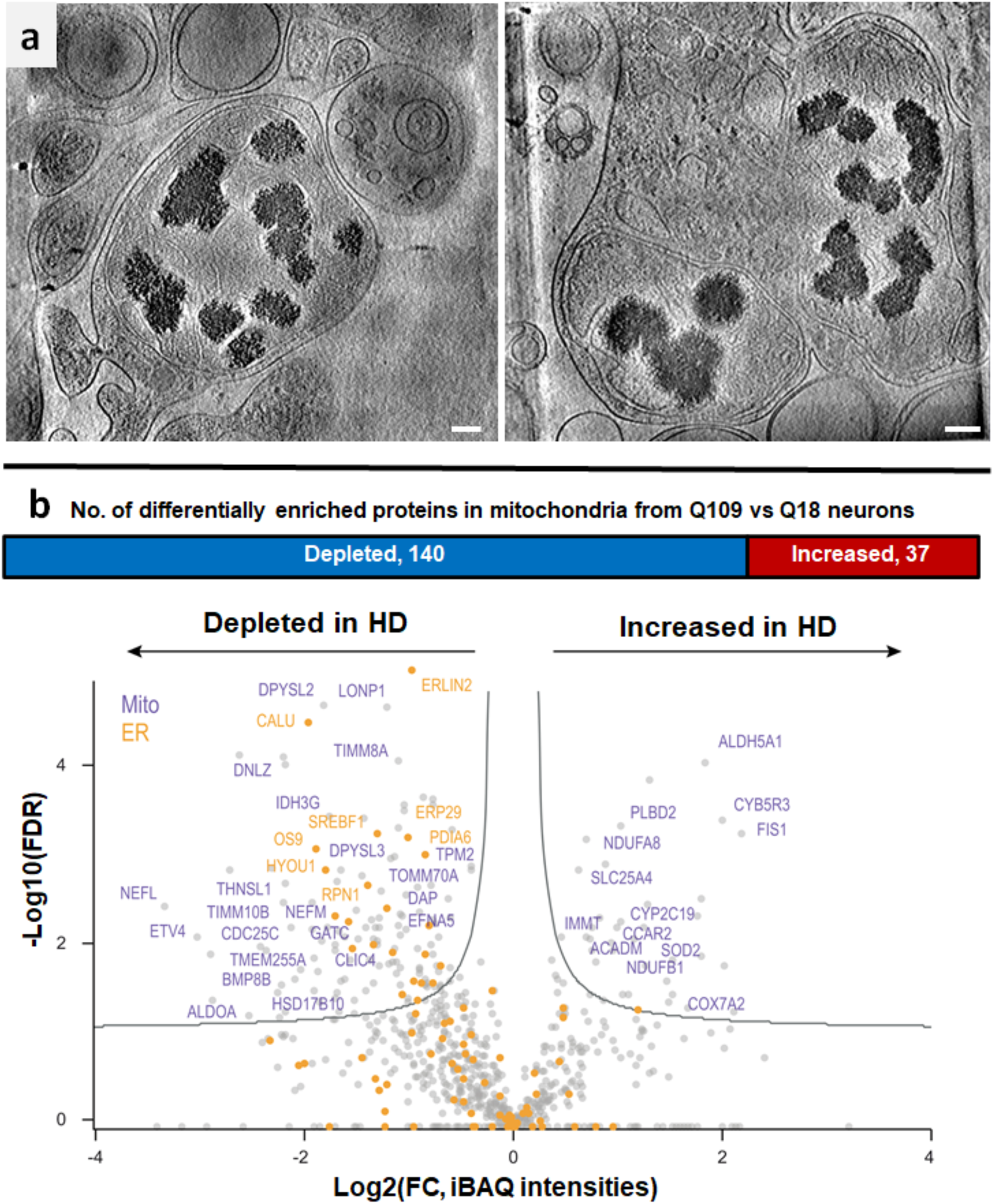
Mass spectrometry of isolated mitochondria revealed differentially enriched proteins (DEPs) in neurites of HD patient iPSC-derived neurons (Q109) vs controls with cryoET showing accumulation of granules in such HD mitochondria. **a.** Z-slices (∼1.4 nm thick) through representative cryoET tomograms of isolated Q109 mitochondria showed abnormal accumulation and enlargement of mitochondrial granules. Scale bars = 100 nm. **b** Mass spectrometry of proteins in isolated mitochondria showed 177 differentially enriched peptides with a potential of 236 protein identities. The scatter plot highlights selected proteins that were depleted or increased in HD (Q109) mitochondria in comparison to controls (Q18). Mitochondrial proteins are highlighted in purple and ER proteins in orange.

Comparing the mitochondrial proteome of Q109 HD patient iPSC-derived neurons vs controls (Q18) revealed a total of 177 differentially enriched peptides (DEPs), with a potential 236 identities, the majority of which were depleted in HD (**Fig 6b** **& Supplementary Table 3**). Differentially enriched proteins included depletion of mitochondrial proteins (purple in **Fig. 6b**) such as TOMM70A, a mitochondrial import receptor involved in the translocation of preproteins into mitochondria, which is impaired in HD^48^. We also detected a depletion of ER proteins in HD mitochondria (orange in **Fig. 6b**), which could reflect impaired mitochondria-ER interactions^57^ and well-documented ER dysfunction in HD^58, 59^. Interestingly, FIS1 (mitochondrial fission protein) levels were increased in HD (**Fig. 6b**), consistent with previous data showing altered regulation of mitochondrial fission in HD^57, 60^.

GO analysis of the differentially enriched peptides found RNA binding to be the most enriched molecular function (**Fig. 7a**), as reflected by increased levels of various cytoplasmic RNA binding proteins such as hnRNPA2B1, hnRNPA1 and hnRNPH1. On the other hand, GRSF1, a mitochondrial RNA binding protein^61^, showed reduced levels in HD mitochondria. This protein is essential for mitochondrial function and required for mitochondrial RNA processing^62^. Indeed, loss of GRSF1 causes mitochondrial stress and can induce senescence phenotypes^63^. Panther pathway analysis identified cytoskeletal regulation by Rho GTPase and pyrimidine metabolism as two other of the three most overrepresented pathways by the DEPs (**Fig. 7b**).

**Fig. 7.**
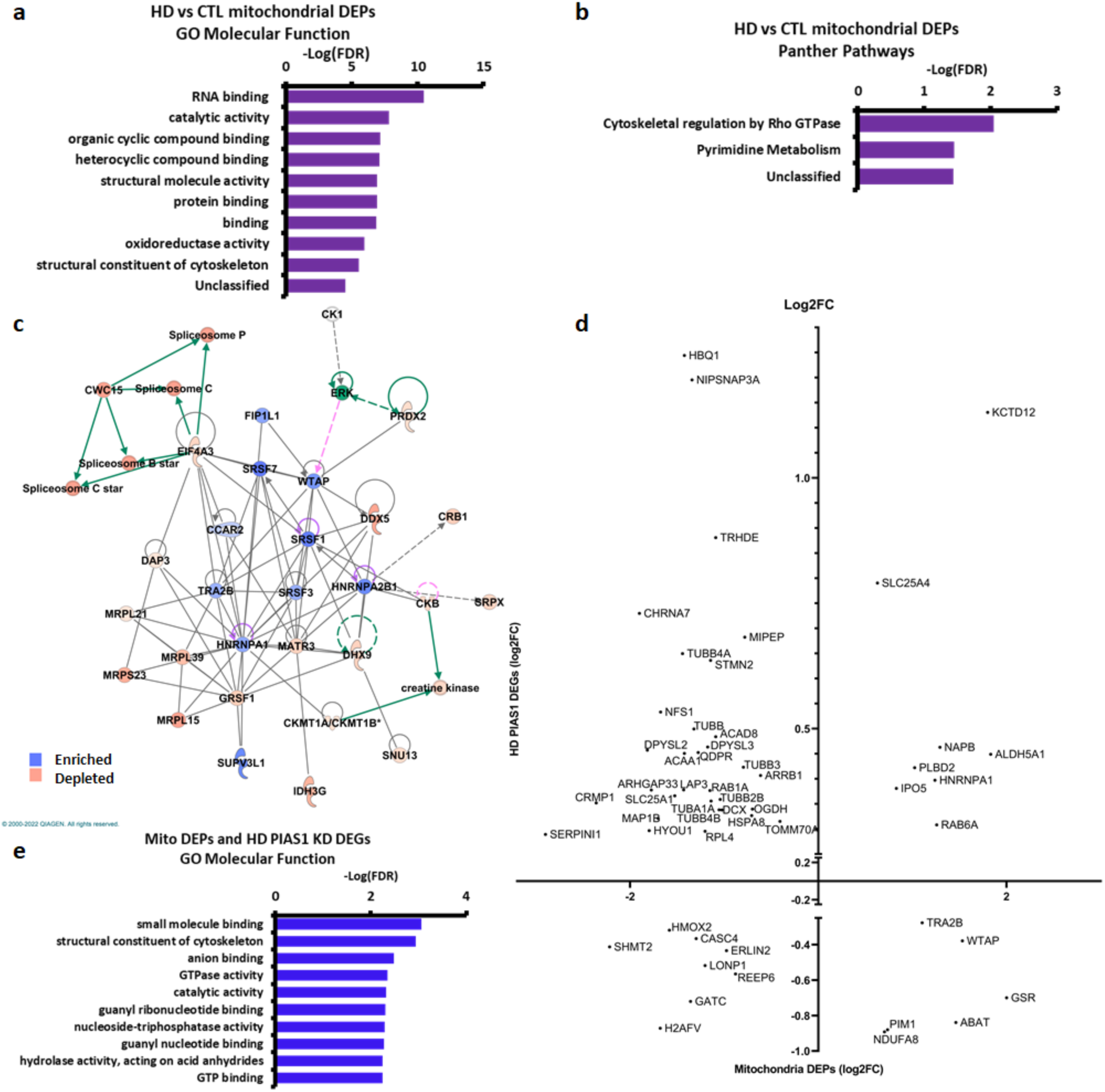
Analysis of mass spectrometry data from isolated HD (Q109) vs control mitochondria showed RNA binding and an overlap with PIAS1 knockdown differentially expressed genes (DEGs). **a** Graph of gene ontology (GO) analysis of the potential 236 identities of the DEPs in HD vs control mitochondria showing molecular functions overrepresented by HD DEPs. **b** Graph of Panther analysis of the 236 DEPs in HD vs control mitochondria showing Panther Pathways overrepresented by HD mitochondria DEPs. **c** Ingenuity pathway analysis of the differentially enriched proteins between HD and control mitochondria highlighted this as the top network (score, 55 Focus molecules: 28), and represents Molecular Transport, RNA Post-Transcriptional Modification, and RNA Trafficking. Proteins in orange are depleted in HD while proteins in blue are enriched. **d** Scatter plot of overlapping DEG log2 fold changes generated from PIAS1 knockdown in HD iPSC-derived neurons from previous work^65^ plotted against the log2 fold enrichment of mitochondrial DEPs. **e** GO analysis of molecular functions overrepresented by the 55 overlapping DEPs/DEGs that are in **d**. N=3 per control and HD samples.

Ingenuity Pathway Analysis (IPA) of the mitochondrial DEPs identified a network representing proteins involved in molecular transport, RNA post-transcriptional modification and RNA trafficking, potentially further implicating these proteins in RNA biology and dynamics (**Fig. 7c**). Upstream regulators identified by IPA included Amyloid Precursor Protein (APP) and transforming growth factor β (TGF-β) as predicted inhibitors of the mitochondrial DEPs in HD. IPA pathways included mitochondrial dysfunction, consistent with the mitochondrial deficits in HD^64^ (**Supplementary Fig. 5c-f**).

### *PIAS1* heterozygous knockout in HD patient iPSC-derived neurons and short-term *Pias1* knockdown in BACHD mouse neurons rescues aberrant mitochondrial granules and presence of sheet aggregates in neurites

Mitochondrial dysfunction can be highly detrimental to neuronal function, particularly in light of the extensive energetic requirements for synaptic function^66^. To evaluate whether the observed phenotypes can be ameliorated, we evaluated genetic reduction of an E3 SUMO ligase, PIAS1, based on our previous data showing that reduced *Pias1* expression resulted in: increased expression of the presynaptic protein synaptophysin in R6/2 HD mice, rescued transcriptional synaptic deficits in zQ175 HD mice, and improved mitochondrial DNA integrity and synaptic gene expression in iPSC-derived neurons^36, 65^. PIAS1 enhances SUMOylation of various proteins, including HTT^45, 67^. To computationally determine whether targeting PIAS1 would predict changes in the HD mitochondrial proteome, we first compared the changes in the mitochondrial proteome of HD iPSC-derived neurons (Q109 vs Q18) described above with findings from our prior study of gene expression in the same type of differentiated neurons following PIAS1 knockdown^65^ (**Fig. 7d**). We found a significant overlap between mitochondrial DEPs and RNA changes induced by siRNA depletion of PIAS1 in HD iPSC-derived neurons (representation factor:1.9 p<2.081e-06, **Fig. 7d**, **Supplementary Fig. 5a**, **Supplementary Table 3**). This overlap of genes and proteins included overrepresentation of GO terms including “GTP binding” and “GTPase activity” as well as several RNA processing-related proteins and tubulins^65^ (**Fig. 7e**). These comparisons further supported investigating whether knockdown of PIAS1 could influence the presence and/or size of the aberrant granules we observed within HD mitochondria.

We again used cryoET to visualize iPSC-derived neurons (Q66, representing an intermediate range of phenotypes) with *PIAS1* heterozygous knockout (hetKO), using a CRISPR-Cas9-generated heterozygous KO, which produces approximately 50% knockdown (**Supplementary Fig. 5e,f**). The *PIAS1* KD iPSC neurons (Q66) differentiated well on EM gold grids in preparation for cryoET experiments (**Supplementary Fig. 5g**). The Q66 *PIAS1* hetKO tomograms (**Fig. 8**) showed seemingly healthy mitochondria and other double membrane-bound organelles (**Fig. 8b**), lacking the abnormally enlarged granules and sheet aggregates, respectively, as had been observed in HD patient iPSC-derived neurons for all Qns (**Fig. 1, 5 & 8a**). Indeed, the structural features of Q66 *PIAS1* hetKO neurites resembled those from control Q18 neurons (**Fig. 8c**) rather than Q66 ones.

**Fig. 8.**
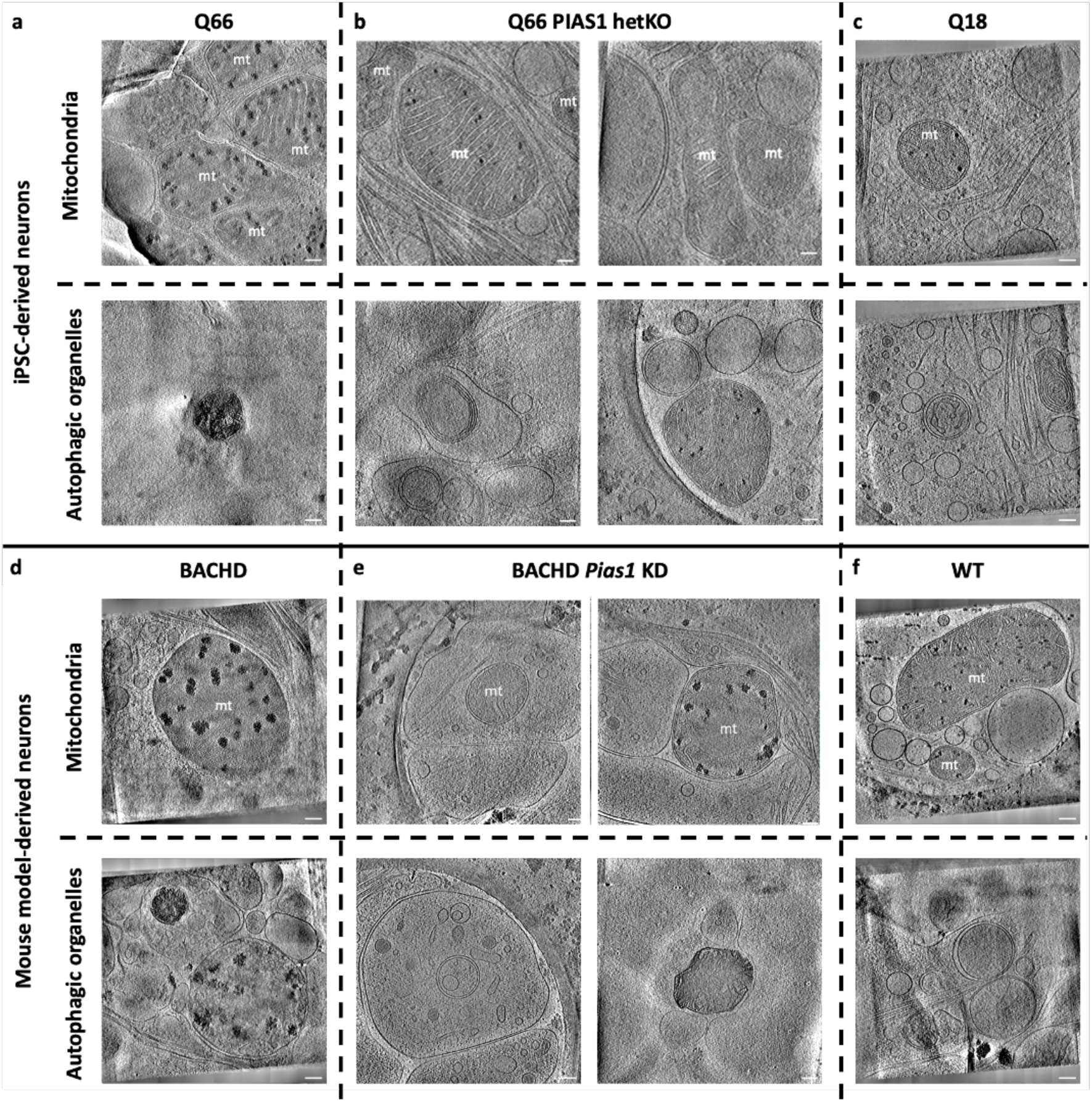
Reduced *PIAS1* rescues mitochondrial granule size and sheet aggregates in neurites of HD model neurons. Slices (∼1.4 nm thick) through representative cryoET tomograms of neurites in **a** Q66, **b** *PIAS1* hetKO, and **c** Q18 iPSC-derived neurons show that *PIAS1* hetKO ameliorated the phenotypes of enlarged mitochondrial granules (first row) and sheet aggregates in putative autophagic organelles (second row) in HD iPSC-derived neurons (Q66). Slices (∼1.4 nm thick) through representative cryoET tomograms of neurites in **d** BACHD, **e** BACHD *Pias1* KD, and **f** WT mouse neurons show that *Pias1* KD ameliorated the enlarged mitochondrial granules (third row) but not the sheet aggregates in putative autophagic organelles (fourth row) in BACHD-derived neurons.

To determine if the rescue of abnormal morphologies observed in HD patient iPSC-derived neurons (Q66) treated with *PIAS1* KD translates to mouse cortical neurons, we carried out a short-term *Pias1* KD in mouse primary neuronal cultures derived from E18 BACHD cortical neurons on EM grids and visualized cells with cryoET (**Fig. 8**). To reduce *Pias1* levels, Accell (Dharmacon) siRNA smart pools against mouse *Pias1* were used. *Pias1* KD was initiated at day *in vitro* 3 (DIV3) with one treatment and grown for 11 days. Cells were vitrified for cryoET analysis at DIV14. Knockdown of *Pias1* in the BACHD neurons was successful according to qRT-PCR analyses; 43% knockdown was achieved comparing control-siRNA-treated neurons to *Pias1* knockdown (**Supplementary Fig. 5i,j**).

CryoET experiments showed that treating BACHD neurons with *PIAS1* siRNA resulted in partial rescue. Indeed, while many mitochondria completely lacked detectable granules, visibly reduced in comparison to the BACHD mitochondria, sheet aggregates in autophagic organelles were present in comparable numbers to those in BACHD neurons without *Pias1* KD (**Fig. 8e**, bottom right). Thus, the beneficial effects of *PIAS1* hetKO in HD patient iPSC-derived neurons was only partially replicated in the mouse model under our experimental conditions, possibly due to the later and shorter treatment time frame. Whether the impact of PIAS1 reduction is consistent across all cells or results in different effects in different cell types needs further investigation.

## Artificial intelligence-based semi-automated 3D segmentation enabled quantification of abnormally enlarged mitochondrial granules in neurites from HD patient iPSC-derived neurons and mouse model primary neurons

We observed both enlarged granules and disrupted cristae in the mitochondria of HD neuronal processes (**Fig. 1-3)**. The mitochondrial granules varied in size, seeming much larger in HD mitochondria than in controls. To quantify their size distribution, we developed a semi-supervised artificial intelligence method to automatically detect and segment mitochondria and mitochondrial granules in the neurite tomograms (**Supplementary Fig. 6**). Among all tomograms (**Supplementary Table 1**), the algorithm found that 139 contained mitochondria for the various HD patient iPSC-derived neurons and 83 for mouse model neurons (**Supplementary Table 1**).

Using quantification of the granule volumes by segmentation estimation, our algorithm revealed that the distribution of mitochondrial granule volumes was shifted towards larger sizes with respect to controls for Q53, Q66, and Q77 (**Figure 9a**), as well as for BACHD, dN17-BACHD, BACHD *Pias1* siRNA, and BACHD transfected with a control siRNA (**Figure 9c**). On the other hand, granule volumes were not larger in Q109 compared to Q18 controls, and were significantly decreased in the Q66 PIAS1 hetKO line when compared to Q66 (Dunn’s multiple comparison Q66 vs. Q66 PIAS1 hetKO padj<0.0001), resembling results for the control Q18 line.

**Fig. 9:**
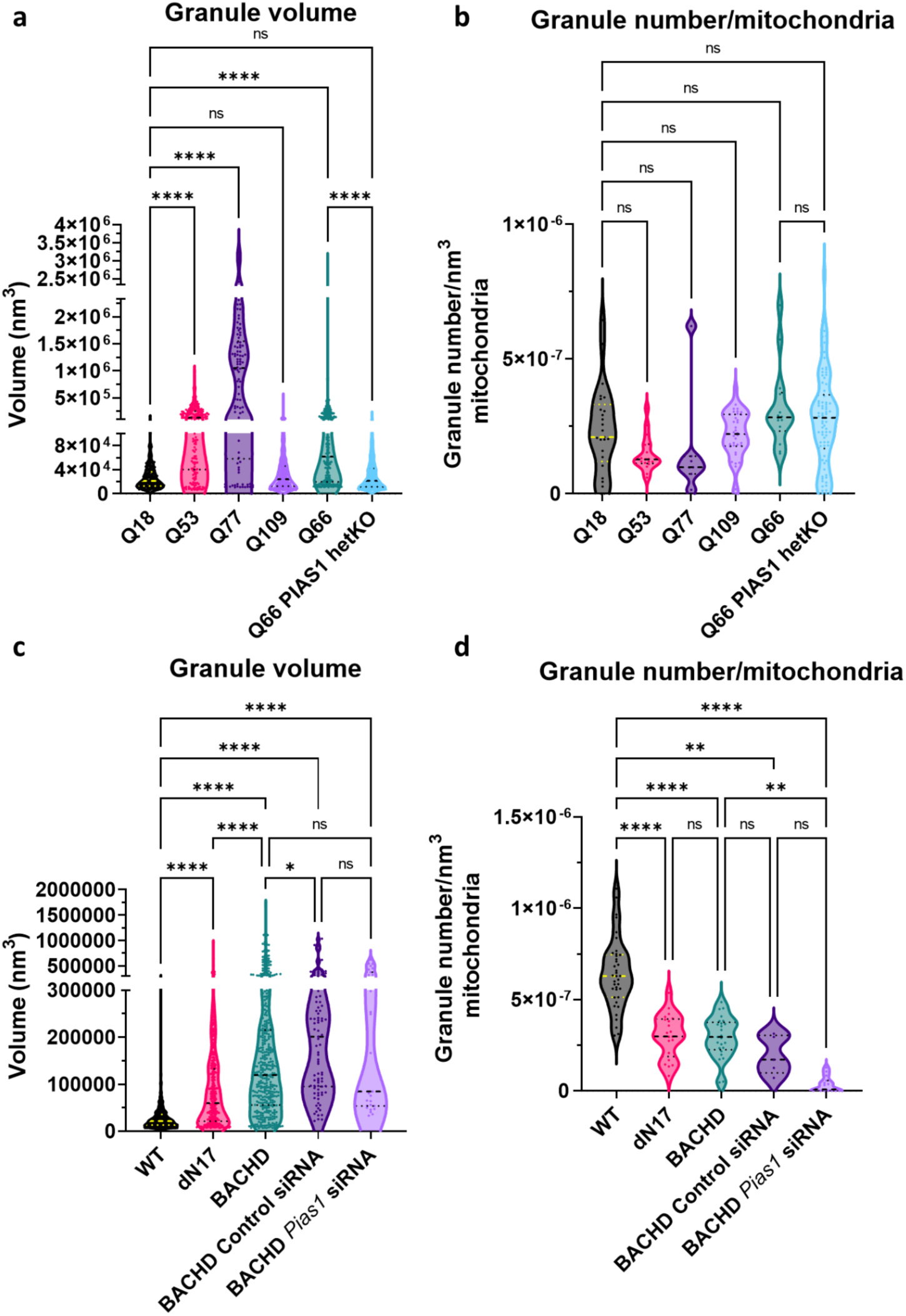
Artificial Intelligence (AI)-based 3D quantification of mitochondrial granule volume and granule number per nm^3^ of mitochondrial volume in cryoET tomograms of mitochondria in neurites demonstrates a significant increase in granule size with higher polyQ in HD human and mouse model neurons and this is rescued in the Q66 PIAS1 hetKO neurons. Violin plots displaying AI measurements of mitochondrial granule **a** volume (Kruskal Wallis statistic = 401.3, P value<0.0001) and **b** numbers per nm^3^ of mitochondrial volume (Kruskal Wallis statistic = 19.78 P= 0.0014) from cryoET tomograms of neurites for five HD patient iPSC-derived neuronal cell lines (tomogram numbers: Q18=21, Q53=14, Q66=10, Q77=5, Q109=37 and *PIAS1* hetKO=68; granule number:Q18=250, Q53=176, Q66=250, Q77=94, Q109=539 and *PIAS1* hetKO=923) as well as mitochondrial granule **c** volume (Kruskal Wallis statistic = 750.8, P<0.0001) and **d** numbers per nm^3^ of mitochondrial volume (Kruskal Wallis statistic = 77.48, P<0.0001) from cryoET tomograms of neurites for three mouse neuron models (tomogram numbers: WT=31, BACHD=22, dN17 BACHD=15, BACHD control siRNA=5 and BACHD *Pias1* siRNA=12; granule numbers: WT=1338, BACHD=380, dN17-BACHD=326, BACHD control siRNA=91 and BACHD *Pias1* siRNA=27). The human neurons and mouse model neurons showed an increase in granule volumes in all but Q109, a trend of reduced granule number in human neurons and a significant reduction in granule number in mouse model neurons. ns=not significant, **** p<0.0001, ** p<0.01, * p<0.05 For full statistical details refer to **Supplementary Table 2**.

Surprisingly, granule volumes were not statistically reduced in BACHD treated with *Pias1* siRNA in comparison to BACHD (Dunn’s multiple comparisons BACHD vs. BACHD *Pias1* siRNA padj>0.9999). On the other hand, control siRNA-treated BACHD neurons showed no difference in granule size distribution compared to *Pias1* siRNA treatment, as expected, given that the latter had no effect (Dunn’s multiple comparisons BACHD Control siRNA vs. BACHD Pias1 siRNA padj>0.9999). Lastly, there was an unexpected modest increase in granule volumes in BACHD upon control siRNA treatment.

While granule volumes were increased for all Qn iPSC-derived neurons except Q109, granule numbers per nm^3^ of mitochondrial volume were not different in any of them from those in control Q18 cells. On the other hand, granule numbers per nm^3^ of mitochondrial volume were decreased in all mouse cell lines with respect to WT, including in those treated with siRNAs.

## DISCUSSION

In this study, we have defined Q-length dependent ultrastructural changes that occur in neuronal processes (neurites) of human HD iPSC-derived neurons and BACHD primary cortical neurons. Specifically, we found that HD neurons contain two types of double membrane-bound organelles with abnormal densities inside, which are completely absent in healthy control neurons. First, we observed ultrastructural changes in neuronal mitochondria, most notably enlarged granules in all HD samples compared to controls (**Fig. 1-3**). Importantly, many HD samples also exhibited severely disrupted cristae, similar to cryoET observations in other neurodegenerative disorders such as Leigh syndrome^68^. Second, we observed sheet aggregates within autophagic organelles resembling mitochondria-derived vesicles, autophagosomes and/or amphisomes (**Fig. 4, 5, Supplementary Fig. 3**). These findings are highly significant in demonstrating the disruption of organellar structure in HD, possibly as very early events in pathogenesis that precede overt neuronal dysfunction and the appearance of inclusions visible in neurons derived from HD patient^21^ and mouse model^20, 69^ brain tissues.

Cellular cryoET experiments ultimately yield 3D volumes (“tomograms”) that sample regions of the crowded subcellular environment in intact cells. Generally, an experienced investigator would use visualization graphics to inspect one tomogram at a time, a laborious discovery process requiring expert knowledge. In the initial stages of this project, we visualized hundreds of neurite tomograms, leading to the discovery of enlarged granules in mitochondria and sheet aggregates in autophagic organelles within them. Using a newly developed semi-supervised artificial intelligence-based method to segment and quantify the number and volume of mitochondrial granules (**Supplementary Fig. 6**), we found that their enlargement is a structural signature of HD, consistently present in both human iPSC- and mouse model-derived neurons (**Fig. 9**).

The aberrant accumulation of large mitochondrial granules and abnormal cristae are known hallmarks of mitochondrial dysfunction as assayed by other methods in similar systems^34, 54^. As members of the HD iPSC Consortium^10^, we previously showed using cell biology techniques that striatal-like HD iPSC-neurons similar to those examined here have mitochondrial deficits including altered mitochondrial oxidative phosphorylation. Indeed, we demonstrated impaired oxygen consumption rate (OCR), altered spare glycolysis capacity (ECAR) and reduced ATP levels in HD neurons^13^. Additional studies have also shown mitochondrial dysfunction, fragmentation and disrupted cristae by traditional electron microscopy of chemically fixed cell lines expressing mHTT^70^.

The high scattering contrast of the enlarged granules we observed here in both human and mouse HD model neurons could be attributable to RNA and/or calcium phosphate, which are more electron dense and thus scatter the electron beam more strongly than most other molecular components in mitochondria comprised of lighter elements such as carbon, nitrogen and oxygen. Our mass spectroscopy data (**Figs. 6, 7 & Supplementary Fig. 4**) are consistent with either interpretation. Indeed, assessing the proteome of mitochondria isolated from human HD iPSC-derived neurons identified differentially enriched proteins related to protein import and RNA binding (**Fig. 7**). RNA granules are normal features of the mitochondrial matrix^61^, and are composed of newly synthesized RNA, RNA processing proteins and mitoribosome assembly factors^61, 71–73^. Stressors such as dysregulation of RNA processing and RNA quality control defects can cause aberrant accumulation of mitochondrial RNA granules, which comprise large ribonucleoprotein complexes^55^. RNA granules are components of mitochondrial post-transcriptional pathways and are responsible for mitochondrial RNA translation^74^. When these granules aberrantly accumulate, the integrity of cristae is compromised to accommodate them^71^. GRSF1, a nuclear encoded RNA-binding protein that regulates RNA processing in mitochondrial RNA granules^62^ and is critical for maintaining mitochondrial function^63, 75^, was significantly decreased in our mitochondrial proteomic dataset in HD. GRSF1 has also been implicated in cellular senescence with levels declining in senescent cells and lowered GRSF1 levels causing mitochondrial stress^63^. Further evidence that these structures may represent mitochondrial RNA granules is the enrichment of RNA binding proteins in the proteome of HD iPSC-derived neurons versus controls.

Alternatively, mitochondria are known to normally contain calcium phosphate granules, which store excess calcium, maintain mitochondrial calcium concentration, and contribute to maintenance of mitochondrial function^76, 77^. Furthermore, calcium overload within cells is known to cause ultrastructural remodeling of cristae^63, 78^, as in our experimental results described above, which was particularly profound for Q109 neurons and dN17-BACHD neurons. Cellular calcium dyshomeostasis is a well-established phenotype in HD^79, 80^, wherein an increase in store-operated calcium entry (SOCE) can lead to increased calcium uptake by the mitochondria due to their proximity to the ER^80^, ultimately resulting in increased mitochondrial granule size due to increased calcium uptake. Furthermore, mitochondrial calcium dysregulation was observed in mitochondria isolated from transgenic YAC128 HD mice^81^. HD mitochondria are more susceptible to calcium stress and form megapores more readily than control mitochondria^82^. Sequestration of calcium into the mitochondria can protect neurons from glutamate excitotoxicity, a phenotype observed in HD neurons^83, 84^. Thus, the enlarged mitochondrial granules we observed here across multiple human and mouse HD model neurons may be composed at least in part of excess calcium phosphate.

The two most likely possibilities for the chemical identity of enlarged mitochondrial granules (*i.e.*, RNA granules versus calcium phosphate granules) need not be mutually exclusive. While future studies such as elemental analysis with electron and/or x-ray microscopy^85^ could help define the chemical nature of these granules, our data provide clues as to their genesis and development.

In iPSCs, granule volume is significantly increased in all cell lines examined herein, except Q109 (**Fig. 9a**), whereas the number of granules per nm^3^ of mitochondrial volume in all iPSC cell lines is not significantly different from that in controls (**Fig. 9b**). This suggests that as new granules or the materials that form them become available, they coalesce with each other and/or with existing ones. On the contrary, for mitochondria in BACHD mice, the larger size of the granules (**Fig. 9c**) is accompanied by a significant reduction in their number (**Fig. 9d**) with respect to controls, suggesting that smaller granules may grow primarily by aggregation with other existing granules rather than by addition of new materials. Further experiments are needed to elucidate the chemical compositions of these granules, which will in turn allow probing for the production of their component materials during neurodegeneration.

Interestingly, granule volumes are larger in iPSC Q66 and Q77 than in our HD mouse models, despite BACHD containing mHTT with a longer polyQ tract (Q97). This may be attributable to differences in the biology of the model systems. Another possible explanation for the statistical discrepancy between cell lines and model systems is that Q53 through Q77 might predictably be in a different (earlier) stage of neurodegeneration than Q109, the latter perhaps being in a state more comparable to BACHD neurons. Importantly, recapitulation in multiple human iPSC lines of the same structural phenotypes seen in mouse primary neurons helps to validate iPSCs as useful HD models.

We also observed differences in the mitochondrial phenotype between the mouse models. While both BACHD and dN17-BACHD mice showed enlarged mitochondrial granules with respect to controls (albeit fewer in number), dN17-BACHD displayed smaller granules than BACHD and severely distorted structures, such as seemingly enlarged mitochondria and swollen cristae. The difference in granule size between BACHD neurons and those treated with control siRNA may be attributable to off-target effects of the control siRNA, timing of treatment or Accell delivery in general. Nonetheless, taken together, our data provide additional hints regarding potential mechanistic underpinnings of normal HTT function. In HD, mitochondrial function appears to be impacted by the altered import of mitochondrial precursor proteins in the presence of mutant HTT^47^. Indeed, the high affinity interaction of mHTT with the inner mitochondrial protein import complex subunit TIM23 disrupts the HD mitochondrial proteome^48^. Furthermore, protein aggregation and alterations in RNA processing and quality control can disrupt protein import^86^. More recently, defects in mitochondrial protein import have also been connected to impaired proteostasis^87^. Mutant HTT can also disrupt mitochondrial trafficking and impair ATP production prior to the appearance of detectable mHTT aggregates^88, 89^. The N17 domain is required for HTT subcellular localization to mitochondria^43, 88, 89^ and the interaction of HTT with the protein import complex TIM23 requires both the N17 and polyQ regions of HTT^47^. Intriguingly, mHTT localizes within the intermembrane space in mitochondria and is more strongly associated with TIM23 than WT HTT, potentially blocking import of nuclear-encoded mitochondrial proteins^48^. This suggests a potential mechanism (**Fig. 10**) underlying the aberrant structures we observe in HD mitochondria (including enlarged granules), particularly for dN17-BACHD, where normal HTT function is impaired by both the lack of the N17 domain and the expansion of the polyQ tract^34,47,54^.

**Fig. 10.**
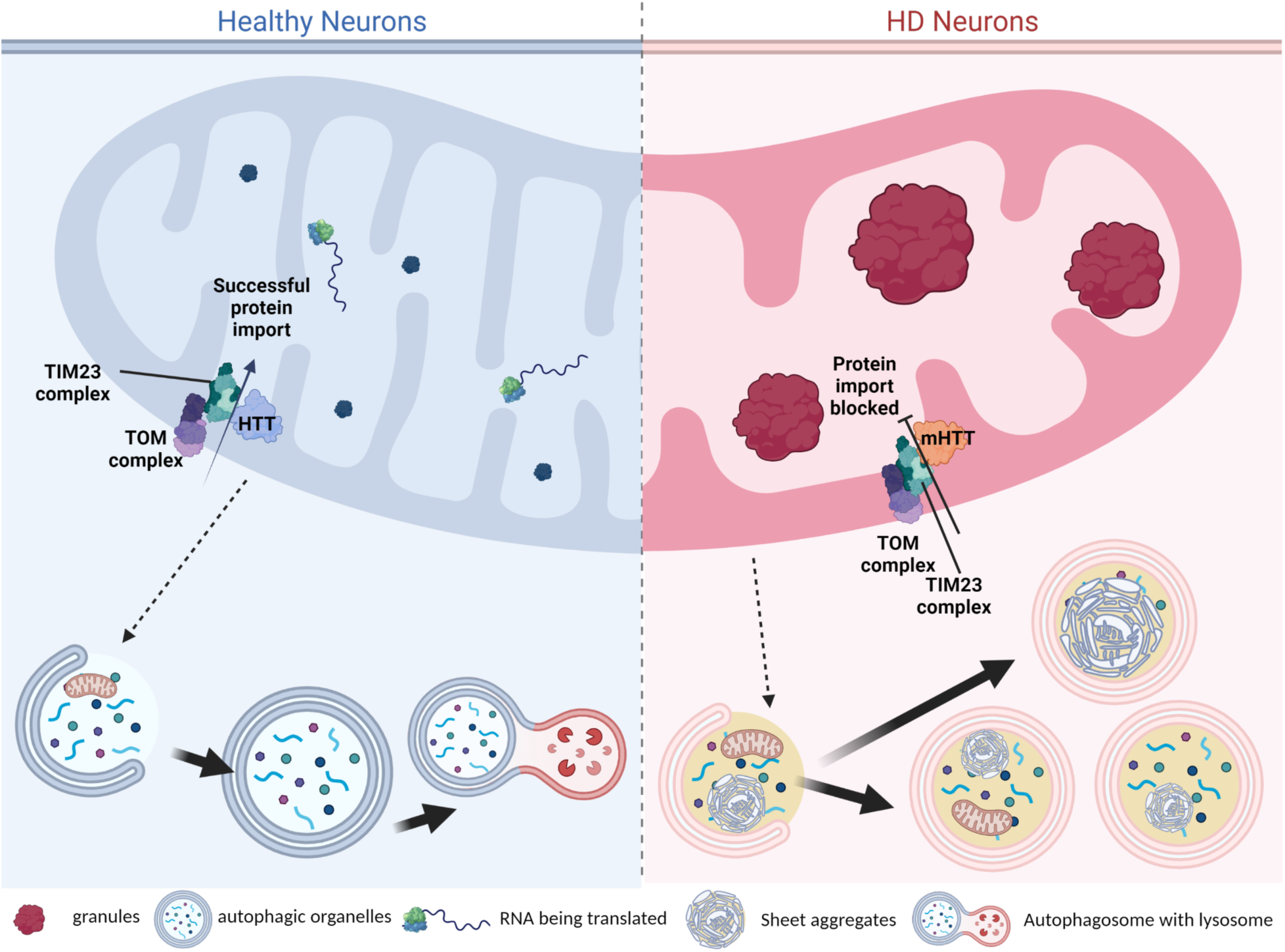
Proposed mechanism of aberrant mitochondrial structures and sheet aggregates in autophagic organelles. Our work here highlights the accumulation of enlarged mitochondrial granules in four human and two mouse HD neuronal models, which we hypothesize could be composed of calcium phosphate and/or enlarged mitochondrial RNA granules that may result from disrupted protein import due to mHTT’s abnormal interaction with the TIM23 complex^48^. Additionally, our mass spectrometry data suggests that RNA binding may be disrupted within HD mitochondria, which may lead to excess RNA molecules within the mitochondrial matrix. There is cross talk between mitochondria, autophagosomes, and lysosomes, and dysfunctional mitochondria are packaged by mitophagy, in addition to other cellular waste; *i.e.*, proteins shown within the autophagic organelles. In healthy neurons, we observe a normal autophagy cascade while in HD we observe accumulation of sheet aggregates in autophagic organelles, which are likely to not be as efficiently processed for degradation as expected in healthy and WT neurons, even though we often see them associated to lysosomes. (Created in BioRender.com).

The sheet aggregates we observed here within autophagic organelles appear to be another early structural hallmark of HD since they were consistently present in the neurites of all HD model systems we examined. In addition to cryoET tomograms, we also collected high-magnification cryoEM images of sample areas with sheet aggregates; the lack of evidence for periodic arrangement in them by Fourier analysis suggests that they are not classical amyloid filaments with the known characteristic cross-beta sheet structures perpendicular to the filament axis^90^; however, this may not be surprising since it is possible that the sheet aggregates we observed may not contain fibrillar mHTT, and HD does not strictly fit within the diseases known as amyloidoses^91^. It is also possible that future purification of the sheet aggregates may render them more amenable to increasingly clarifying high-resolution analyses without interference from the cell membrane and surrounding milieu of the crowded cytosol. Whether the sheet aggregates contain mHTT also awaits future biochemical analyses of purified autophagic organelles and/or degenerated mitochondrial particles. Nevertheless, our data here suggest that these unusual structural features, likely in components of autophagic pathways, are a consequence of the stress and toxicity induced by the presence of full-length, endogenous mHTT or fragments derived from it.

Given the potential hypotheses suggested by the observation that the sheet aggregates may be degenerated mitochondria (**Supplementary Fig. 3e,f**) associated with lysosomes (**Supplementary Fig. 3g**), it would be expected that instances of mitochondria that simultaneously exhibit some recognizable cristate, cristae junctions, and granules, at the same time as some clear sheet aggregate densities, might constitute extremely rare events that are hard to capture along a dynamic degeneration process. That is, the sheet aggregates may develop from remodeled and compromised mitochondrial membranes and/or granules through a previously unresolved mechanism in HD cells endogenously expressing untagged mHTT, and may go through intermediate states where mitochondrial features are extremely challenging to recognize or no longer present due to degeneration, before the overt appearance of sheet aggregate densities within them. Indeed, we observed many double membrane-bound compartments in most of our specimens whose identities were not clearly assignable but that may actually correspond to mitochondria in these hypothesized intermediate states, such as the organelle in the bottom left of **Supplementary Fig. 3e,** and those in **Supplementary Fig. 3a-d**. In support of this hypothesis, mitochondria have been recently demonstrated to interact directly with lysosomes via their membranes^53^. Indeed, the traditional paradigm for largely separate functions by these two organelles has changed with emerging evidence that mitochondria and lysosomes are mutually functional and interdependent, with profound implications to the underpinnings of aging^92^ and neurodegeneration^93, 94^.

In spite of the challenges, this exciting observation provides an additional hint into resolving the potential origin of the sheet aggregates. Mitochondrial autophagy (mitophagy) is a clearance mechanism of defective mitochondria via autophagy^35^ and is altered in many neurodegenerative disorders such as Parkinson disease (PD) and Alzheimer disease (AD)^95^. There is now emerging evidence that mitophagy is also defective in HD. Altered mitophagy may also potentially contribute to the bioenergetic deficits observed in various HD models^13, 96^ and at least in part to the excessive weight loss that is characteristic of late-stage human HD^97, 98^. Consistent with our findings, recent work^31^ has identified non-fibrillar mHTT within single-membrane-bound organelles in cortical and striatal tissue from zQ175 HD mice, including multivesicular bodies (MVB) and amphisomes, using correlative light and conventional electron microscopy of samples fixed by freeze substitution. The localization of non-fibrillar mHTT changed depending on the disease stage, with presymptomatic stages showing localization within MVBs/amphisomes and late stage disease showing localization to the autolysosomes or residual bodies^31^. Our findings here in intact, cryo-preserved human patient and mouse model-derived neurons are suggestive of even earlier events and thus are likely complementary.

Our previous studies showed improved synaptic-associated gene expression and mitochondrial DNA integrity in HD iPSC-neurons after PIAS1 KD^36, 65^. Based on these data, we evaluated the effect of reduced PIAS1 levels on the mitochondrial and autophagosome structural phenotypes in HD patient iPSC- and mouse model-derived neurons and observed rescue of these phenotypes in the former and partial rescue in the latter (**Fig. 8** **& 9**). Interestingly, our findings on PIAS1 KD also inform on the possible biochemical composition of mitochondrial granules. For example, we previously identified MCUR1, CALM1, CALB1 and CABP1 as differentially expressed in iPSC-derived neurons upon *PIAS1* knockdown^65^, consistent with changes to calcium phosphate granules. We also observed alterations in the levels of RNA binding and processing genes, consistent with changes to mitochondrial RNA granules. Several mitochondrial proteins are SUMO modified (identified in yeast) and require Siz1 or Siz2 for modification (*i.e.*, PIAS1/2)^99^. For instance, Fis1, involved in mitophagy, is SUMO modified and reduction of SUMO modification of Fis1 restores appropriate mitophagy in Hela cells^100^. Finally, a recent report identified PIAS1 as a potential age-of-onset modifier. PIAS1 containing a S510G single nucleotide polymorphism, which reduces SUMO modification of mHTT^101^, delays HD onset and produces milder disease severity in HD mice, consistent with the concept that reduced PIAS1 levels may ameliorate disease. Our studies thus represent a proof of concept that the synergistic combination of cryoET and proteomics of iPSC- and mouse model-derived neurons or organelles within them can inform on the impact of a given therapeutic strategy on structural features and ultimately function, and may be applicable to other cellular systems and disease models.

## METHODS

### Ethics Statement and mouse models

All protocols involving the use of animals in the study, namely BACHD^32^ and deltaN17 BACHD^33^, were approved by the Institutional Animal Care and Use Committee at the University of California in San Diego.

### iPSC culture, differentiation, and maintenance

Neuronal differentiation was performed once iPSC colonies reached 60-70% confluency as previously described^4^. Differentiation was initiated by washing iPSC colonies with phosphate buffered saline pH 7.4 (PBS – Gibco) and switching to SLI medium (Advanced DMEM/F12 (1:1) supplemented with 2 mM Glutamax^TM^ (Gibco), 2% B27 without vitamin A (Life technologies), 10 µM SB431542, 1 µM LDN 193189 (both Stem Cell Technologies), 1.5 µM IWR1 (Tocris)) with daily medium changes. This was day 0 of differentiation; at day 4, cells were pretreated with 10 µM Y27632 dihydrochloride (Tocris) and then washed with PBS and then passaged 1:2 with StemPro Accutase (Invitrogen) for 5 minutes at 37 °C and replated onto plates which were coated with hESC qualified matrigel (1 hour at 37 °C) in SLI medium containing 10 µM Y27632 dihydrochloride for 1 day after plating and continued daily feeding with SLI medium until day 8. At day 8, cells were passaged 1:2 as above and replated in LIA medium (Advanced DMEM/F12 (1:1) supplemented with 2 mM Glutamax^TM^, 2% B27 without vitamin A, 0.2 µM LDN 193189, 1.5 µM IWR1, 20 ng/ml Activin A (Peprotech)) with 10 µM Y27632 dihydrochloride for 1 day after plating and daily feeding was continued through day 16. At day 16, cells were plated on either the carbon side of Quantifoil holey carbon film grids (Electron Microscopy Sciences; see below for grid preparation for cryoET), or in 6 well Nunclon coated plates with PDL and hESC matrigel for mitochondrial isolation. Cells were plated at 1x10^6^ for mitochondrial isolation and at half density for cryoET of neurites in intact neurons at 500,000 per dish on the 35 mm Mat-tek glass bottomed dish containing 3 holey-carbon grids in SCM1 medium (Advanced DMEM/F12 (1:1) supplemented with 2 mM Glutamax^TM^, 2% B27 (Invitrogen), 10 µM DAPT, 10 µM Forskolin, 300 µM GABA, 3 µM CHIR99021, 2 µM PD 0332991 (all Tocris), to 1.8 mM CaCl2, 200 µM ascorbic acid (Sigma-Aldrich), 10 ng/ml BDNF (Peprotech)). Cells on EM grids were topped up with an additional 1 ml of SCM1 (35 mm Mat-Tek dishes). We tried even lower densities to improve the probability of getting one cell per grid square; however, the neurons did not survive or did not differentiate well. Medium was 50% changed every 2-3 days. On day 23, medium was fully changed to SCM2 medium (Advanced DMEM/F12 (1:1): Neurobasal A (Gibco) (50:50) supplemented with 2 mM Glutamax^TM^), 2% B27, to 1.8 mM CaCl2, 3 µM CHIR99021, 2 µM PD 0332991, 200 µM ascorbic acid, 10 ng/ml BDNF) and 50% medium changes were subsequently performed every 2-3 days. Cells were considered mature and ready for subsequent analyses and experiments at day 37.

### Mouse model neuronal culture and maintenance

Established protocols were followed to set up cortical neurons collected from mouse E18 embryos ^102–104^. Briefly, cortical tissues were extracted from E18 mouse embryos and extensively rinsed in HBSS with 1% Penicillin-Streptomycin, followed by dissociation in 0.25% trypsin with 1 mg/ml DNase I. Cortical neurons were isolated and plated with plating media (Neurobasal with 10% FBS, 1xB27,1xGlutaMAX) onto either glass coverslips for immunostaining or into 12 well plates for biochemistry at appropriate density. Both the cover glasses and plates were pre-coated with poly-L-lysine (Invitrogen). Plating medium was replaced with a maintenance medium (Neurobasal, 1xB27, 1xGlutaMAX) the following day. Only 2/3 of the media was replaced every other day until the conclusion of the experiments.

### Mitochondria isolation

Day 37 neurons were harvested using the Miltenyi MACS human mitochondria isolation kit (Miltenyi Biotec 130-094-532) with additional proteases added to the lysis buffer aprotinin (10 μg/ml), leupeptin (10 μg/ml), PMSF (1 mMl), EDTA-free protease inhibitor cocktail III (1x SIgma-Aldrich - 539134) at a concentration of 10 million/ml of lysis buffer and disruption was performed using an 27 G needle using a syringe to triturate 10 times up and down and then proceeding to labeling following the manufacturer’s recommended protocol. Final mitochondrial pellet was resuspended in 50 μl of storage buffer and flash frozen in liquid nitrogen, stored at -80 °C until mass spec analysis.

### RNA extraction & concentration

RNA extraction was performed using QIAGEN RNEasy kit following manufacturer’s protocol. RNA was eluted in 50 μl of nuclease free H2O and required further concentration for cDNA synthesis. RNA concentration was performed using ZYMO RNA Clean and Concentrator^TM^-5 kit.

### cDNA synthesis

100 ng of RNA was used for cDNA synthesis using the Quantabio qScript® cDNA SuperMix kit while 25 ng of the RNA was used for RT-reaction of just RNA and nuclease-free water. cDNA synthesis was performed using the manufacturer’s protocol at 25 °C for 5 minutes, then 42 °C for 30 minutes, and finally 85 °C for 5 minutes in a Bio-Rad T100 thermal cycler.

### qRT-PCR

One μl of cDNA at a concentration of 5 ng/μl was used per reaction, with 10 μM primers for mouse *Pias1* and *Eif4a2* for a housekeeping control using previously published primers^65^. Quantification was performed on a Quantstudio 5 using SYBR green reagent (Biorad) running delta delta CT method and calculating fold change normalizing to non-transgenic primary neurons control SMARTPool siRNA. Prism 9.0 was used to calculate statistical significance by two way ANOVA.

### Western blot

Protein was harvested from frozen cell pellets (for CRISPR validation of the clone) using RIPA lysis buffer or from isolated mitochondria samples described above. Samples were subjected to SDS (sodium dodecyl sulfate) polyacrylamide gel electrophoresis and Western blotting onto nitrocellulose. Membranes were assessed using either infrared fluorescence on the Li-Cor. Antibodies were as follows: LC3ab (Cell Signaling #12741), PIAS1 (Cell Signaling #3550), CTIP2 (abcam #ab18465), ATPB (abcam #ab14730), normalization of loading was calculated based on REVERT total loading stain.

### Immunofluorescence

Immunofluorescence staining on day 37 neurons was performed as previously described^4^

### Pias1 knockdown in neurons

Knockdown of *Pias1* was performed for mouse model neurons using Accell SMARTPool siRNA against *Pias1* (Horizon Discovery cat#E-059344-00-0005) and a non-targeting SMARTPool control (Horizon Discovery cat#D-001910-10-05) for comparison. Treatment was performed at 3 days *in vitro* (DIV3) at a concentration of 1 μM in 1 ml of medium per 35 mm MatTek dish, medium was topped up to 2 ml after 24 hours and then 2/3 of the media was replaced every other day until the conclusion of the experiments.

### CRISPR-Cas9 heterozygous knockout of PIAS1 in iPSC

Clones were selected from stem cell edited pools in a method that was previously described^65^

### Grid preparation for cryoET of HD patient iPSC-derived neurons and mouse model neurons

For iPSCs, Quantifoi®l R 2/2 Micromachined Holey Carbon grid: 200 mesh gold (SPI supplies Cat#:4420G-XA) grids were prepared for cell plating by sterilizing using forceps to carefully submerge them in 100% ethanol (Fisher Scientific) at an angle perpendicular to the liquid surface and then passed quickly through a yellow flame. Grids were then placed into 1 ml of poly-D-lysine (100 µg/ml in borate buffer, pH 8.4) in a 35 mm Mat-Tek glass coverslip bottom dish (VWR P35G-0-14-C) at an angle perpendicular to the liquid’s surface, and coated on the bottom of the dish for at least 1 hour at room temperature. When cells were ready for plating, the grids were washed two times with sterile-filtered Milli-Q H2O and a final wash with PBS before cell plating.

For mouse neurons, EM grids were briefly dipped into 70% ETOH to sterilize, followed by coating with 0.1mg/ml PDL^105^. The grids were rinsed with sterile dH2O and loaded with isolated neurons. The maintenance of these grids were exactly as described above. DIV14 neurons were used for cryoEM/cryoET experiments.

For both iPSC-derived and mouse model neurons, cells grown on grids were vitrified using the temperature- and humidity-controlled Leica GP2. Grids were retrieved from culture dishes using the forceps for the Leica device, and 3 µl of 15 nm BSA gold tracer (Fisher Scientific; Catalog No.50-248-07) was pipetted onto the carbon- and cell-side of the grid. Grids were loaded and blotted for 5 seconds in 95% humidity at 37 °C and immediately plunged into liquid propane. The vitrified grids were transferred into grid storage boxes in clean liquid nitrogen and then stored in clean liquid nitrogen prior to cryoET imaging.

Cells on the remainder of the dishes that were used for cryoET, were scraped using a cell scraper in cold PBS and pelleted at 350 xg for 5 minutes, PBS was aspirated and the pellets were flash-frozen in liquid nitrogen and stored at -80 °C for later analysis of knockdown.

### CryoEM/ET data collection

For each specimen, namely vitrified HD patient iPSC-derived and mouse primary neurons, we collected low magnification (6500X) cryogenic TEM images and assembled them into montages to screen for regions of interest (ROI) before cryoET tilt series collection. All images were acquired using a G3 Titan Krios^TM^ cryo-electron microscope (ThermoFisher Scientific) operated at 300 kV, energy filter at 30 eV, in low-dose mode using SerialEM software^106^. At each tilt angle, we recorded “movies’’ with 5-6 frames per movie using a Gatan K2^TM^ or K3^TM^ direct electron detection camera with a BioQuantum^TM^ Imaging Filter (Gatan, Inc). The tilt series were collected at higher magnification (39000X, 3.47 Å/pixel sampling size) using a dose-symmetric tilting scheme^107^ from 0°, target defocus of −5 μm and a cumulative dose of ∼120 e/Å^2^.

### Tomographic reconstruction

Upon data collection, all tilt series were automatically transferred to our computing and storage data clusters and images were motion-corrected using MotionCor2^108^ and reconstructed into full tomograms automatically using EMAN2^109^. This on-the-fly reconstruction facilitated the screening of tomograms.

After screening the automated cryoET reconstructions, we used IMOD^110^ software for standard weighted-back projection tomographic reconstruction of tomograms with interesting and relevant features, using coarse cross-correlation-based alignment, gold fiducial-based alignment, or patch tracking alignment. For each tilt series, unsuitable images with large drift, excessive ice contamination, etc. were manually removed prior to tilt series alignment. For cryoET tilt series containing prominent sheet aggregates and subjected to Fourier analysis and subtomogram averaging attempts to determine whether they contained repeating features, we corrected for the contrast transfer function (CTF) using IMOD’s recent 3D-CTF correction algorithm and reconstructed using a SIRT-like filter (16 iterations)(**Fig. 4** and **Supplementary Fig. 3**).

### Tomographic annotation and segmentation

The tomograms containing mitochondrial granules were post-processed binning by 4x and applying various filters, such as a low-pass Gaussian filter at a frequency=0.01, a Gaussian high-pass filter to dampen the first 1-5 Fourier pixels, normalization, and thresholding at 3 standard deviations away from the mean. For both types of tomograms (containing mitochondrial granules or sheet aggregates), we carried out tomographic annotation of different features in binned-by-4 tomograms using the EMAN2 semi-automated 2D neural network-based annotation^42^, and performed manual clean-up of false positives in UCSF Chimera^111^. The cleaned-up annotations were thresholded, low-pass filtered, and turned into binary masks, which were multiplied by contrast-reversed versions of the tomograms to segment out each corresponding feature.

### Visualization of tomograms containing sheet aggregates

Initially, the sheet aggregates appeared filamentous in 2D z-slices through our cryoET tomograms. When attempts at subtomogram averaging failed to yield averages with filamentous morphology, and instead resulted in featureless sheets, we more carefully examined the tomograms slice-by-slice to understand the causes for this apparent failure. This revealed that the linear features in a section persisted through multiple sections above or below. While this might be expected due to lower resolvability in z because of the missing wedge, we also noticed that the linear features shifted their location from section to section (as if drifting sideways), and in different directions. This anomaly suggested that the linear features seen in 2D slices corresponded to sheets in 3D, oriented at various angles with respect to the x-y plane, and explained the preliminary exploratory subtomogram averages that had also revealed a sheet morphology.

The sheets seem to be composed of extremely electron dense materials yielding high contrast in cryoEM/ET images, and are exceedingly thin (∼20 Å, a bit more or less in some regions), as measured from 3D-CTF corrected tomograms without any downsampling or low-pass filtration.

### Quantification and statistical analysis of mitochondria granules size

We developed a two-stage deep learning system for voxel level annotation of mitochondria and granules in tomograms. In the first stage, our system is trained on a handful of annotated slices on a subset of tomograms and learns to segment mitochondria and granules. In the second stage, we use our model to make predictions on all the tomograms and use high confidence predictions as pseudo-annotations to augment our training set. We then train a new model on this augmented training set and use its predictions to quantify the number and sizes of mitochondria and granules in the tomograms.

In the first stage, we train a 3D-UNet^112^ model to perform segmentation of the 3D volumes containing objects of interest. We train two separate models, one for segmenting mitochondria and the other for segmenting granules. These models are trained in a semi-supervised fashion on sparsely annotated 3D volumes - only 2% of the 2D slices are manually annotated with each pixel being labeled as being part of the background or being part of the mitochondria / granule.

Since annotating each slice is a time-consuming process, we utilize pseudo-labelling to generate more annotations. In the second stage, we run every tomogram through our model and add the high confidence predictions to the training set. Next, we retrain our model on the expanded training set which consists of both human- and machine (pseudo)-labeled slices. The new model is then used to make the final voxel level predictions on all the tomograms.

We refine the 3D segmentations in the post-processing stage. We run a connected components analysis to count the number of segmented objects. We filter out background noise in the predictions by discarding objects which don’t fall within the expected range of volumes. We retain granule detections which are located within a detected mitochondrion. After post-processing, we scale each voxel by a factor of 21.02 to get the volume in nm^3^.

Compilation of the data into graphs was performed in Prism 9.3 (https://www.graphpad.com/scientific-software/prism/) with statistics performed based on the number of groups to compare and data normality. iPSC-derived neurons used Dunn’s multiple comparisons to compare control to the various HD lines, for the mouse primary neurons, Dunn’s multiple comparisons were used to compare WT vs all other groups and then BACHD vs dN17 and Control siRNA vs *Pias1* siRNA treatment.

### Sample preparation for proteomic analysis

Isolated mitochondria were solubilized in a final concentration of 1% SDS and mitochondrial proteome was extracted using methanol-chloroform precipitation. 400 µl methanol, 100 µl chloroform and 350 µl water were added sequentially to each 100 µl sample, followed by centrifugation at 14,000 x g for 5 min at room temperature. The top phase was removed and the protein interphase was precipitated by addition of 400 µl methanol, followed by centrifugation at 14,000 g for 5 min at room temperature. Pellet was air dried and resuspended in 8M urea, 25 mM ammonium bicarbonate (pH 7.5). Protein concentration was determined by BCA (Thermo Fisher) and 2-4 µg total protein were subjected to reduction and alkylation by incubation with 5 mM DTT for 1 h at room temperature followed by 5 mM iodoacetamide for 45 min at room temperature, in the dark. The samples were then incubated with 1:50 enzyme to protein ratio of sequencing-grade trypsin (Promega) overnight at 37 °C. Peptides were acidified with trifluoroacetic acid to a final concentration of 1%, desalted with μC18 Ziptips (Millipore Sigma), dried and resuspended in 10 μL 0.1% formic acid in water.

### LC-MS/MS acquisition

LC-MS/MS analyses were conducted using a QExactive Plus Orbitrap (QE) mass spectrometer (Thermo Fisher) coupled online to a nanoAcquity UPLC system (Waters Corporation) through an EASY-Spray nanoESI ion source (Thermo Fisher). Peptides were loaded onto an EASY-Spray column (75 μm x 15 cm column packed with 3 μm, 100 Å PepMap C18 resin) at 2% B (0.1% formic acid in acetonitrile) for 20 min at a flow rate of 600nl/min. Peptides were separated at 400 nL/min using a gradient from 2% to 25% B over 48 min (QE) followed by a second gradient from 25% to 37% B over 8 minutes and then a column wash at 75% B and reequilibration at 2% B. Precursor scans were acquired in the Orbitrap analyzer (350-1500 m/z, resolution: 70,000@200 m/z, AGC target: 3x10^6^). The top 10 most intense, doubly charged or higher ions were isolated (4 m/z isolation window), subjected to high-energy collisional dissociation (25 NCE), and the product ions measured in the Orbitrap analyzer (17,500@200 m/z, AGC target: 5e4).

### Mass spectrometry data processing

Raw MS data were processed using MaxQuant version 1.6.7.0 (Cox and Mann, 2008). MS/MS spectra searches were performed using the Andromeda search engine^113^ against the forward and reverse human and mouse Uniprot databases (downloaded August 28, 2017 and November 25, 2020, respectively). Cysteine carbamidomethylation was chosen as fixed modification and methionine oxidation and N-terminal acetylation as variable modifications. Parent peptides and fragment ions were searched with maximal mass deviation of 6 and 20 ppm, respectively. Mass recalibration was performed with a window of 20 ppm. Maximum allowed false discovery rate (FDR) was <0.01 at both the peptide and protein levels, based on a standard target-decoy database approach. The “calculate peak properties” and “match between runs” options were enabled.

All statistical l tests were performed with Perseus version 1.6.7.0 using either ProteinGroups or Peptides output tables from MaxQuant. Potential contaminants, proteins identified in the reverse dataset and proteins only identified by site were filtered out. Intensity-based absolute quantification (iBAQ) was used to estimate absolute protein abundance. Two-sided Student’s *t*-test with a permutation-based FDR of 0.01 and S0 of 0.1 with 250 randomizations was used to determine statistically significant differences between grouped replicates. Categorical annotation was based on Gene Ontology Biological Process (GOBP), Molecular Function (GOMF) and Cellular Component (GOCC), as well as protein complex assembly by CORUM.

Additional analysis was performed on all potential identities of the differentially enriched/depleted proteins that were significant by t-test, using Panther pathways and Panther Overrepresentation algorithms for GO Molecular Function, GO Biological Processes and GO Cellular Component at http://www.pantherdb.org/. Ingenuity Pathway Analysis was performed using the significantly differential proteins to assess pathways, networks and upstream regulators. For comparison of DEPs to PIAS1 knockdown DEGs, overlap statistics for overrepresentation was performed at http://nemates.org/MA/progs/overlap_stats.html using total genome number of genes at 20,500.

## ACKNOWLEDGEMENTS

We thank Dr. William X. Yang’s group at University of California in Los Angeles for providing the BACHD and deltaN17-BACHD mouse models, Weijiang Zhou for feedback on analysis of high-resolution images of sheet aggregates to look for possible repeating features, and Ronald Courville for assistance with generating manual labels to train the artificial intelligence algorithm to find and quantify mitochondrial granules. We thank the support of NIH grants (P01NS092525 to LMT, WC, JF, WM; S10OD021600 to WC; R35NS116872 to LMT), postdoctoral fellowships from the Hereditary Disease Foundation to GHW and CSG, and Chan Zuckerberg Initiative Neurodegeneration Challenge Pairs Pilot Project to WC and SY (2020-221724, 5022).

## AUTHOR CONTRIBUTIONS

G-H.W., C.S-G., J.G.G-M., P.M., L.M.J., M.F.S, C.W., W.M., J.F., L.M.T., & W.C. were involved in conception and design of the experiments related to differentiation and cryo electron tomography.

G-H.W., J.G.G-M., M.F.S., &W.C. analyzed cryo-ET data.

C.S-G., optimized iPSC cell growth on EM grids, and performed all iPSC differentiations and cell culture.

Y.G., performed all mouse model neuronal cultures.

G-H.W., L.M.J., & P.M. optimized cryoET grid preparation and screening and collected all cryoET data.

J.G.G-M. & G-H.W. performed tilt series alignment and tomographic reconstruction. J.G.G-M., G-H.W., and C.D. performed tomographic annotation.

J.G.G-M., G-H.W., & M.F.S. participated in cryoET data visualization.

C.S-G, R.A., J.F., and L.M.T. were involved in conception and design of the experiments for the mitochondria mass spectrometry.

C.S-G, R.A., N.R.G., and L.M.T. were involved in the acquisition, the validation, the analysis and interpretation of the mitochondrial mass spectrometry data.

C.S-G., Y.G., R.M., and K.Q.W. were involved in the differentiation and cell culture of the neurons and validation of PIAS1 knockdown.

S.G., J.H. & S.Y were involved in creating an artificial intelligence-based algorithm for automated segmentation of features in cryoET tomograms and quantitative analyses of mitochondrial granules.

G-H.W., J.G.G-M., & C.D. were involved in extensive manual reference labeling to train artificial intelligence algorithms for segmentation and quantification of features in cryoET tomograms.

G-H.W, C.S-G, J.G.G-M., R.A., C.W., S.G., S.Y., L.M.T. & W.C, wrote the manuscript.

G-H.W, C.S-G, J.G.G-M., M.F.S., L.M.T. & W.C., substantively revised the manuscript with input from all authors.

## DATA AVAILABILITY

We will deposit representative tomograms for each phenotype in EMDB (https://www.emdataresource.org/deposit.html); and in Chorus, for proteomics.

