## Supplementary Information for "CryoET Reveals Organelle Phenotypes in Huntington Disease Patient iPSC-Derived and Mouse Primary Neurons"

### SUPPLEMENTARY FIGURES AND TABLES

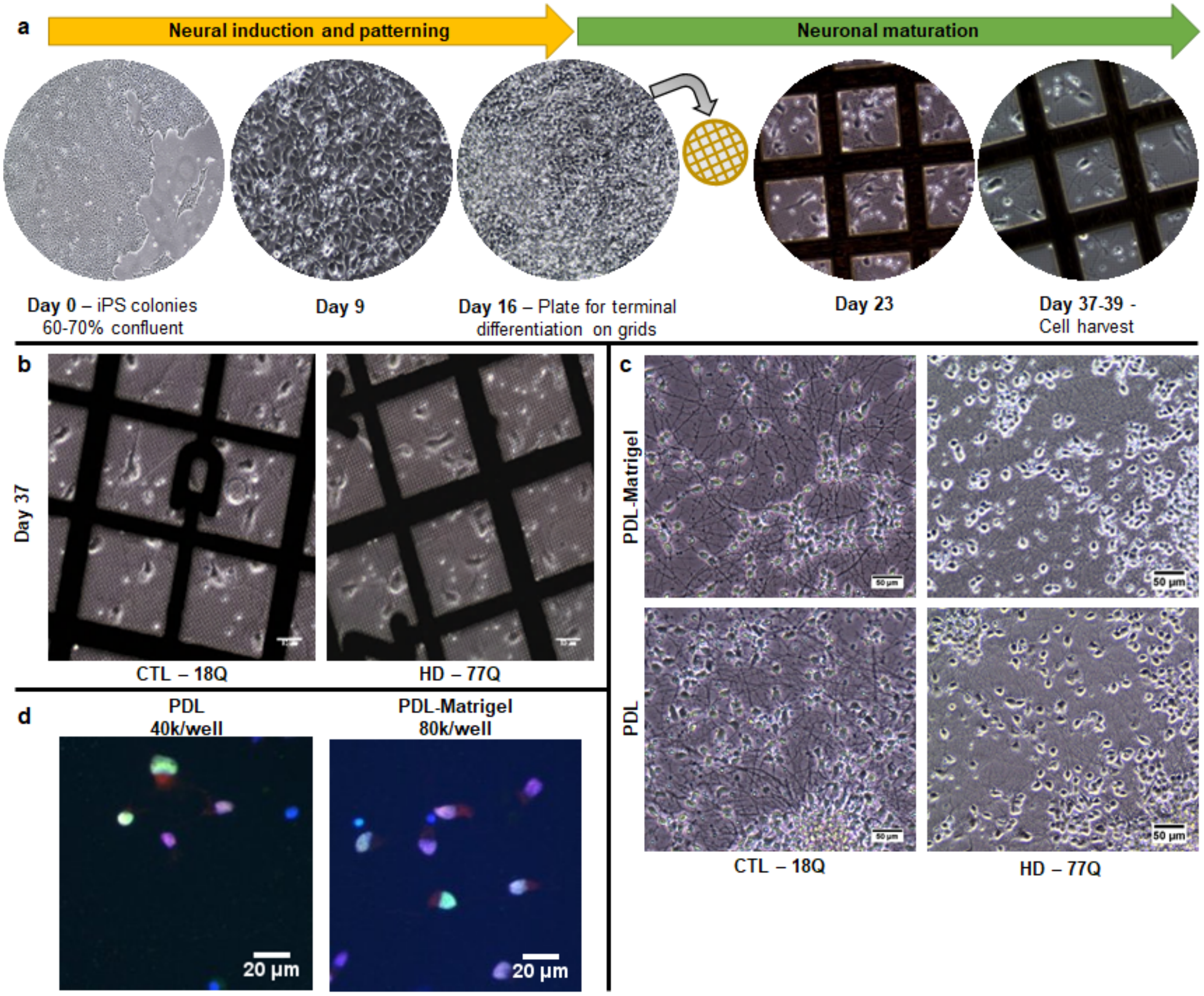


**Supplementary Fig. 1. Differentiation of control and HD iPSC-derived neurons for cryoET. a** Differentiation paradigm outlining our previously published general protocol[4], introducing here the modification of plating cells at day 16 onto carbon grids for cryoET. **b** Representative phase-contrast images of day 37 neurons that were grown on grids coated with PDL alone, adapted from published protocol. Scale bar = 15 µm. **c** Representative images of control and HD Day 37 iPSC-derived neurons differentiated as previously described on PDL and Matrigel and compared to just PDL alone. Scale bar = 50 µm. **d** Immunofluorescence of the 53Q line for two MSN markers, CTIP2 in green and DARPP32 in red, at our experimental setup for the grids on the left and cultured as previously described[4] on the right. Scale bar = 20 µm.

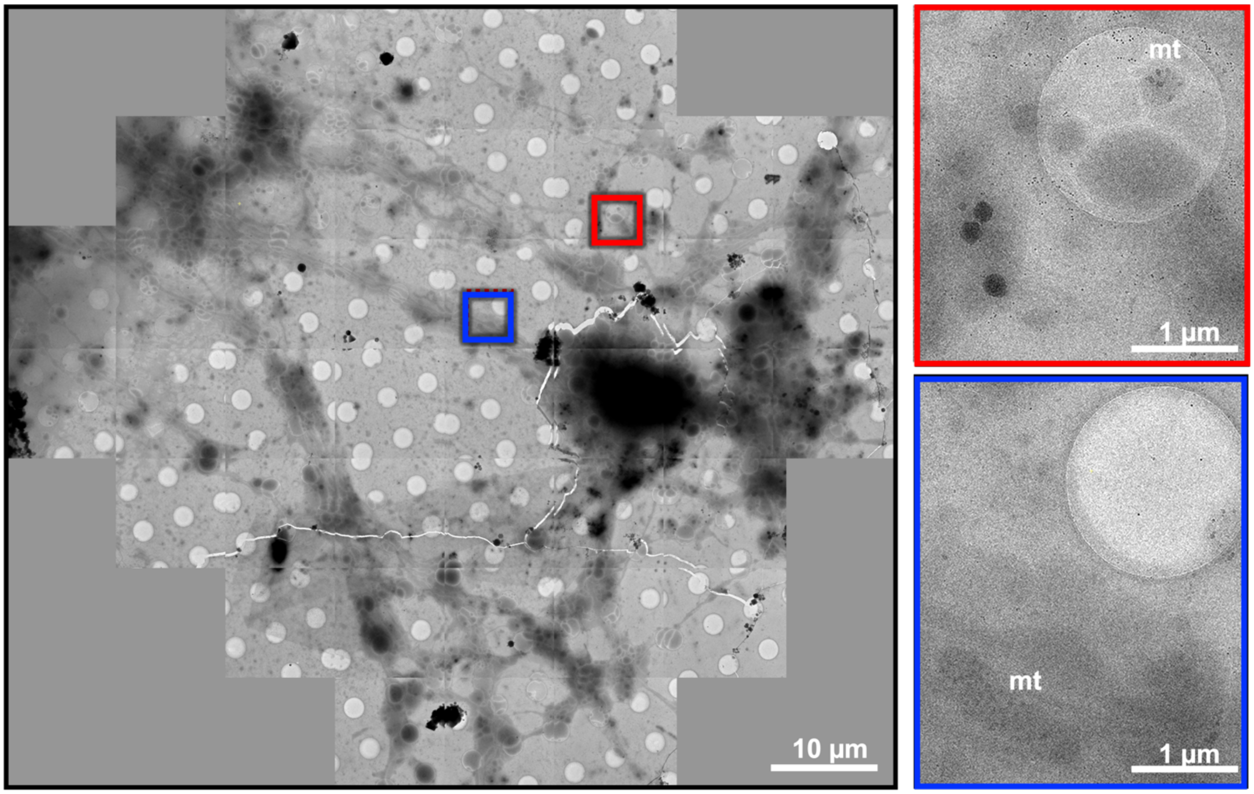


**Supplementary Fig. 2. HD patient iPSC-derived neurons grow well on cryoEM grids.** A montage of low-magnification (6500 X) images (n=56) of HD patient iPSC-derived neuron cryoEM images (left), and intermediate magnification (39000 X) screening images (right) from the regions highlighted with a red box in the montage, showing putative mitochondria (labeled as mt) with visibly enlarged and dense granules inside. Scale bars = 10 **μ**m and 1 **μ**m.

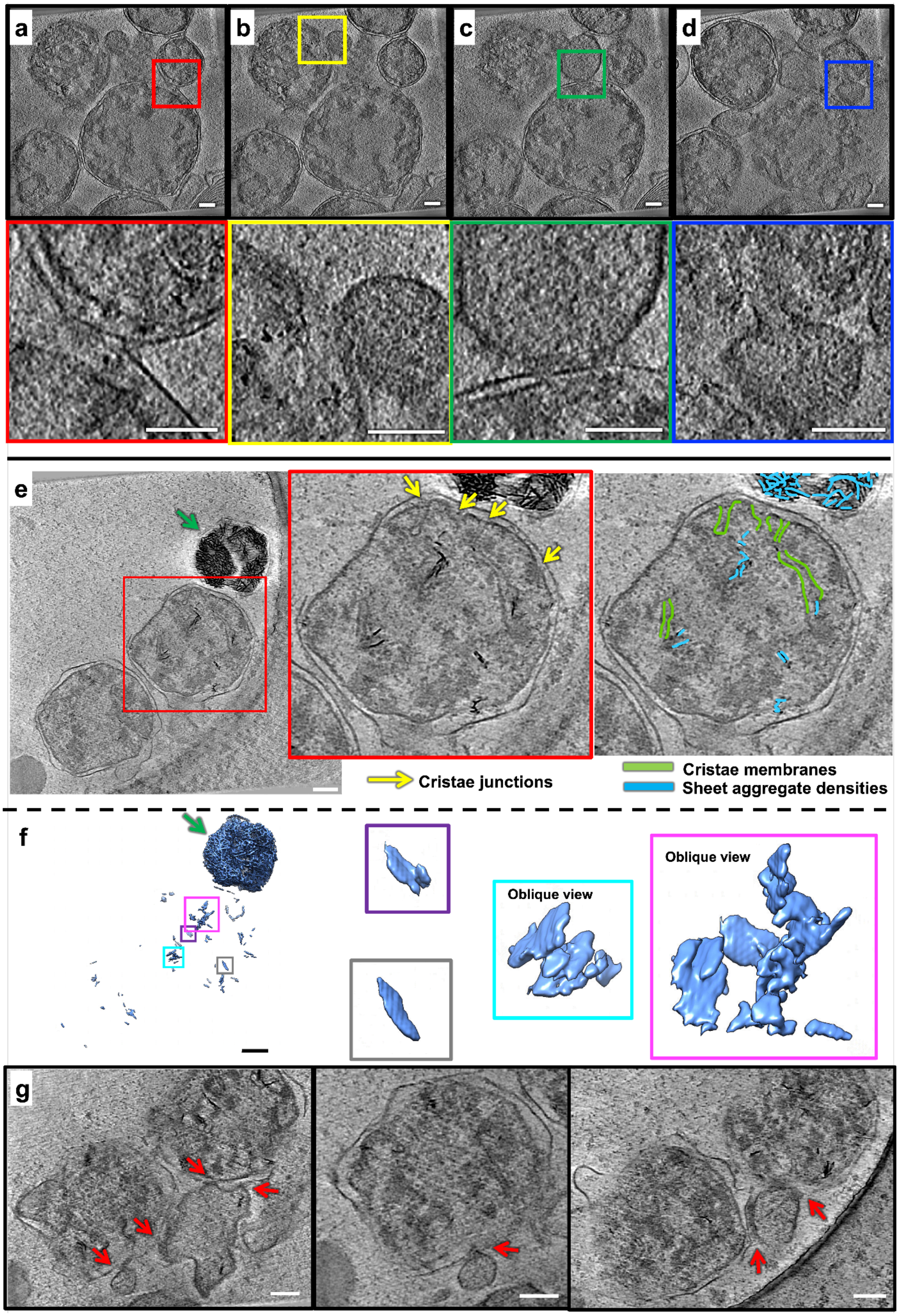
a
b
c
d
e
Cristae junctions
Cristae membranes
Sheet aggregate densities
f
Oblique view
Oblique view
g

**Supplementary Fig. 3. Aggregates inside double membrane-bound compartments in a neurite of HD patient iPSC-derived neuron (Q53).** Slices (~14 nm thick) through selected regions of a representative cryoET tomogram showing aggregates in double membrane-bound compartments (putatively organelles in the autophagic pathway) in a neurite of an HD patient iPSC-derived neuron (Q53), with blown-up views highlighting the compartments potentially fusing with **a** each other or **b-d** with single membrane-bound compartments (putatively lysosomes). **e** Slices (~1.4 nm thick) through another tomogram (Q53) showing 3 double membrane bound compartments, with the one in the top right of the image showing a compartment completely overwhelmed by sheet aggregates, and the one in the middle showing incipient sheet aggregate densities and structural hallmarks of mitochondria, such as a double membrane, cristae, and cristae junctions. Semi-automated, neural-net based annotation with EMAN2 of sheet aggregate densities, training on a few positive references (n=10) from the sheet aggregate pointed at with the green arrow in the top right, identifies densities in the other membrane-bound compartments automatically as belonging to the same feature as the well-recognized, mature sheet aggregate in the top, free from bias. The blown-up and oblique (pink and cyan boxes) views on the right clearly show the sheet-like morphology of the incipient densities in the mitochondria-like compartment in the middle. **g** Tomographic slices (~56 nm thick) showing that the mitochondria-like organelle in **e** and its neighbor to the bottom-left are interacting with single membrane-bound compartments (at sites indicated by the red arrows), putatively lysosomes, similar to the double-membrane bound organelles shown in **a-d**. Scale bars are 100 nm.

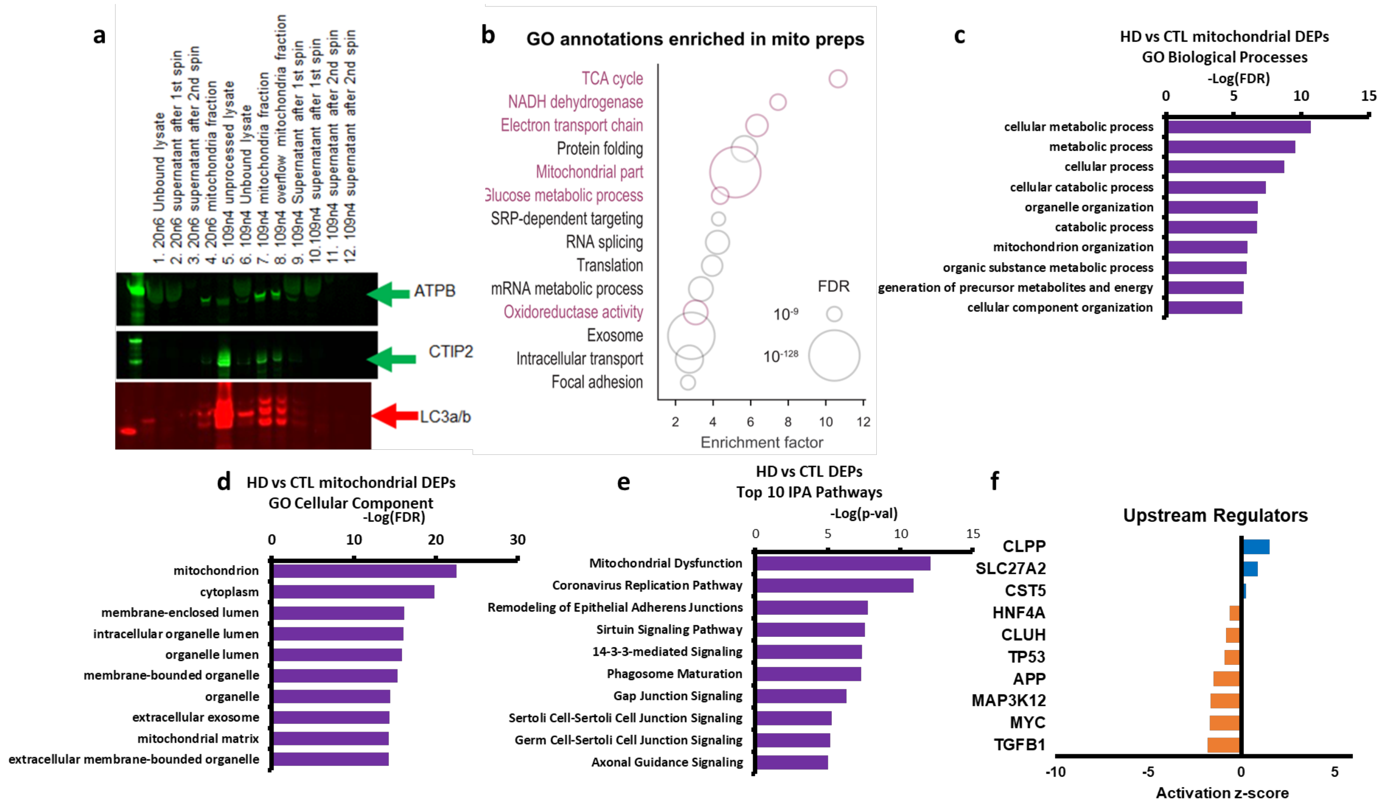


**Supplementary Fig. 4. Mass spectrometry of isolated mitochondria. a** Representative Western blot on various fractions from the enrichment of mitochondria on a pilot experiment using MACS-based isolation probing for mitochondrial markers. **b** GO annotations show enrichment of mitochondrial proteins in isolated mitochondria from control and HD neurons. **c-f** IPA and GO analyses of HD vs control neurons highlighting significant DEPs that show the overrepresentation of proteins related to **c** GO biological processes and **d** GO cellular components. **e** Top 10 IPA pathways by p value and **f** Top 10 IPA Upstream regulators (genes and proteins) by p value that have assigned activation scores. **Supplementary Table 3** shows full GO and IPA lists.

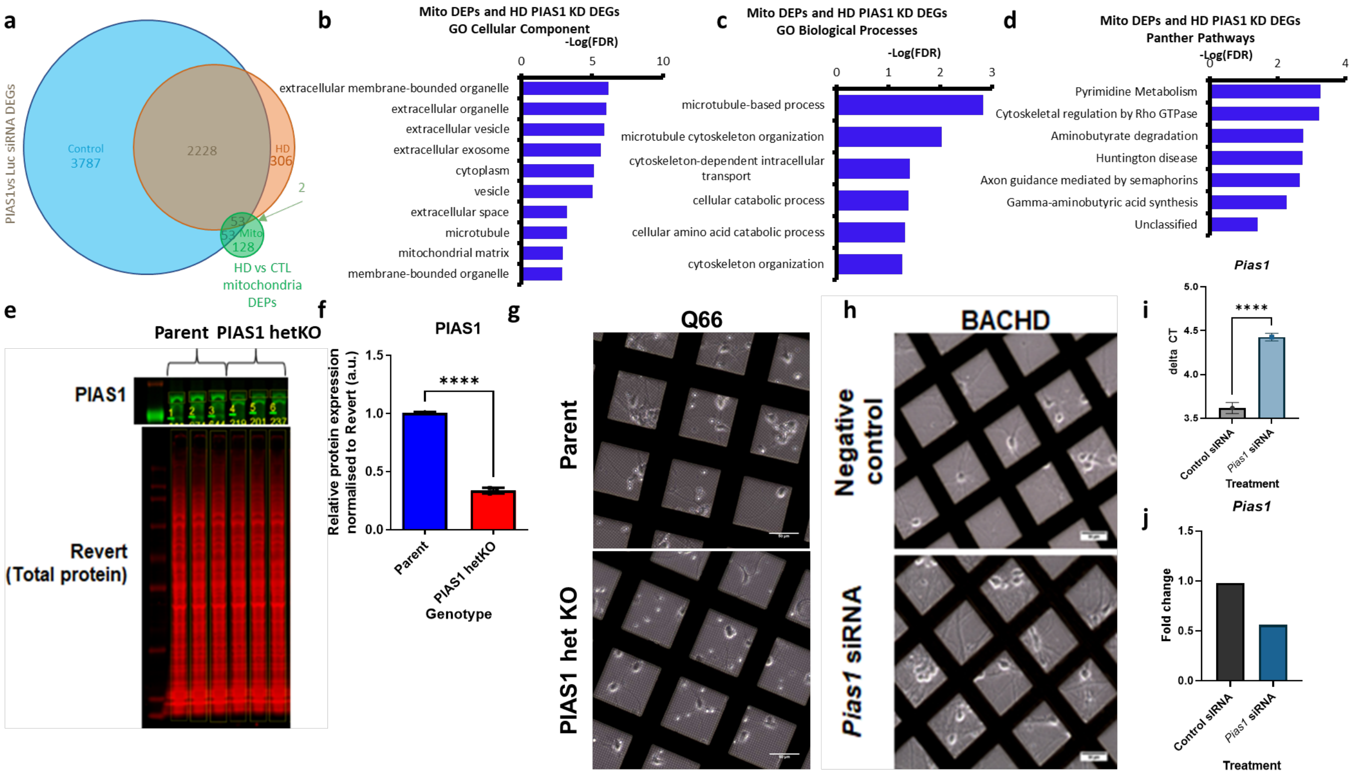


**Supplementary Fig. 5. Knockdown of PIAS1 to rescue effects observed by cryoET.** **a** Venn diagram that shows the total PIAS1 knockdown generated DEGs in control and HD iPSC-derived neurons and the overlap with the HD vs control mitochondrial DEPs. **b-d** Assessing the 55 common genes between PIAS1 knockdown in HD neurons and HD mitochondria show overrepresentation of these terms for **b** GO Cellular component, **c** GO Biological processes and **d** Panther Pathways. **Supplementary Table 3** shows full GO terms lists. **e** Western blot of the iPSC that were the parental and CRISPR edited line showing PIAS1 knockdown and this is quantified in **f** (Unpaired two-tailed t-test t=46.70, df=4, p<0.0001). **g** Representative images of the iPSC derived neurons at day 37 prior to vitrification for cryoET showing PIAS1 knockdown does not affect cellular growth on the grids **h-j** E18 BACHD neurons at DIV14, Pias1 siRNA treatment was performed at DIV3, **h** Representative images of the primary neurons growing on grids prior to vitrification, **i-j** qRT-PCR for Pias1 to validate Pias1 knockdown in BACHD primary neurons, graphs show **i** delta CT (Unpaired two-tailed t-test t=18.23, df=4 p<0.0001) and **j** Fold change showing significant knockdown.

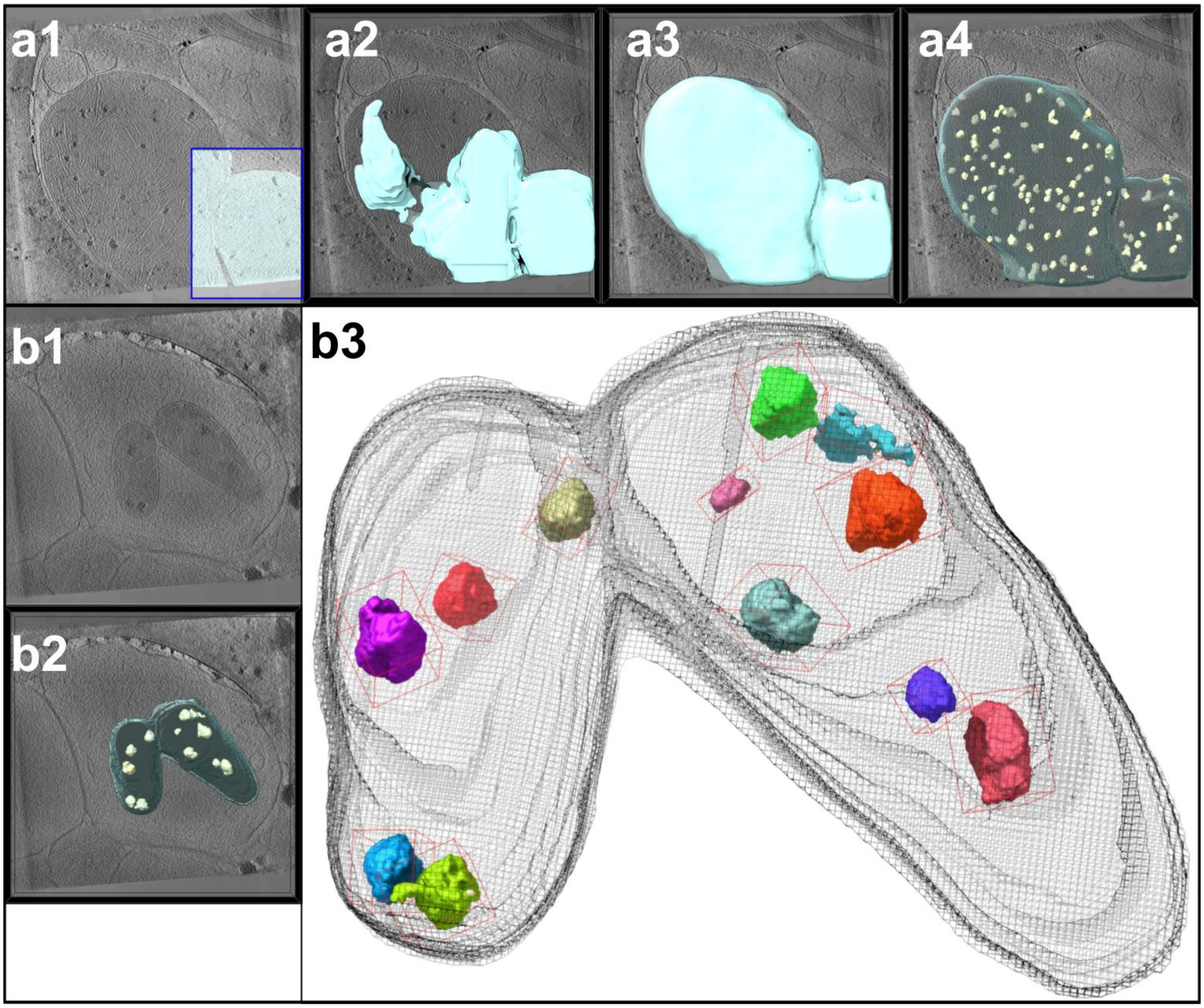


**Supplementary Fig. 6. Granule and Mitochondria Segmentation Pipeline. a1** - The 3D UNet for mitochondrial segmentation is trained on a handful of partially annotated slices. Pixels in the blue rectangle are labeled as being part of the mitochondria or the background and the rest are unlabelled. **a2** - High confidence predictions (shown in cyan) from the 3D UNet are used as pseudo-labels to augment the training set. A new 3D UNet is trained on the augmented dataset. **a3** - Retraining the 3D UNet on this augmented dataset improves segmentation quality. The new segmentation (shown in cyan) spans the full extent of the mitochondria. **a4** - The 3D UNet used to detect granules is applied to the mitochondrial volume. The resulting predictions are shown in yellow. **b1** - A previously unseen tomogram is fed into the trained 3D UNets. **b2** - The mitochondria and granule predictions produced by the segmentation pipeline. **b3** - A connected components analysis is used to identify individual granules and measure their volumes.

**Supplementary Table 1.** Number of EM grids in iPSC-differentiation or mouse primary neuron petri dishes and tilt series collected from them, reconstructed into tomograms. "M" in the second to last column indicates the number of tomograms containing mitochondria with visible granules used for quantification in **Fig. 9** while "A" indicates the number of tomograms containing large sheet aggregates.

| | | | | | |
|---|---|---|---|---|---|
| **iPSC** | **Q18** | No. of EM grids | 3 | **M** | 21 |
| | | No. of final tomograms | 46 | **A** | 4 |
| | **Q53** | No. of EM grids | 3 | **M** | 14 |
| | | No. of final tomograms | 46 | **A** | 5 |
| | **Q66** | No. of EM grids | 3 | **M** | 10 |
| | | No. of final tomograms | 31 | **A** | 8 |
| | **Q66 PIAS1 KD** | No. of EM grids | 3 | **M** | 68 |
| | | No. of final tomograms | 71 | **A** | 3 |
| | **Q77** | No. of EM grids | 3 | **M** | 5 |
| | | No. of final tomograms | 10 | **A** | 6 |
| | **Q109** | No. of EM grids | 3 | **M** | 37 |
| | | No. of final tomograms | 42 | **A** | 6 |
| **Mouse** | **WT** | No. of EM grids | 3 | **M** | 31 |
| | | No. of final tomograms | 34 | **A** | 3 |
| | **BACHD** | No.of EM grids | 3 | **M** | 22 |
| | | No. of final tomograms | 24 | **A** | 14 |
| | **BACHD RNAi control** | No. of EM grids | 3 | **M** | 5 |
| | | No. of final tomograms | 29 | **A** | 3 |
| | **BACHD PIAS1 KD** | No. of EM grids | 3 | **M** | 12 |
| | | No. of final tomograms | 36 | **A** | 5 |
| | **dN17-BACHD** | No. of EM grids | 3 | **M** | 15 |
| | | No. of final tomograms | 34 | **A** | 6 |

**Supplementary Table 2**: Kruskal-Wallis statistics that pertain to data displayed in **Fig. 9**

| **Sample** | **Measurement** | **Kruskal-Wallis summary** | **Multiple comparison test** | **Multiple comparisons stats** |
|---|---|---|---|---|
| Human iPSC-neurons | Granule volume | K-W Stat = 401.3 No. of groups = 6, P value<0.0001, no. of values = 2222 | Dunn's | Q18 vs. Q53 padj<0.0001 |
| | | | | Q18 vs. Q66 padj<0.0001 |
| | | | | Q18 vs. Q77 padj<0.0001 |

| | | | | |
|---|---|---|---|---|
| | | | | Q18 vs. Q109 padj=0.9211 |
| | | | | Q18 vs. Q66 PIAS1 hetKO padj>0.9999 |
| | | | | Q66 vs. Q66 PIAS1 hetKO padj<0.0001 |
| Mouse primary neurons | Granule volume | K-W Stat = 750.8 No. of groups = 5 P value<0.0001, no. of values = 2162 | Dunn's | WT vs. BACHD padj<0.0001 |
| | | | | WT vs. dN17-BACHD padj<0.0001 |
| | | | | WT vs. BACHD Control siRNA padj<0.0001 |
| | | | | WT vs. BACHD *Pias1* siRNA padj<0.0001 |
| | | | | BACHD Control siRNA vs. BACHD *Pias1* siRNA padj>0.9999 |
| | | | | BACHD vs. BACHD Control siRNA padj=0.0189 |
| | | | | BACHD vs. BACHD *Pias1* siRNA padj>0.9999 |
| | | | | BACHD vs. dN17-BACHD padj<0.0001 |
| Human iPSC-neurons | Granule count/$nm^3$ of mitochondria | K-W Stat = 19.78 No. of groups = 6, P value = 0.0014, no. of values = 177 | Dunn's | Q18 vs. Q53 padj=0.1913 |
| | | | | Q18 vs. Q66 padj=0.7658 |
| | | | | Q18 vs. Q77 padj=0.5736 |
| | | | | Q18 vs. Q109 padj>0.9999 |
| | | | | Q18 vs. Q66 PIAS1 hetKO padj>0.9999 |
| | | | | Q66 vs. Q66 PIAS1 hetKO padj>0.9999 |
| Mouse primary neurons | Granule count/$nm^3$ of mitochondria | K-W Stat = 77.48 No. of groups = 5 P value<0.0001, no. of values = 104 | Dunn's | WT vs. BACHD padj<0.0001 |
| | | | | WT vs. dN17-BACHD padj<0.0001 |
| | | | | WT vs. BACHD Control siRNA padj=0.0013 |
| | | | | WT vs. BACHD *Pias1* siRNA padj<0.0001 |
| | | | | BACHD Control siRNA vs. BACHD *Pias1* siRNA padj>0.9999 |
| | | | | BACHD vs. BACHD Control siRNA padj>0.9999 |

| | | | | |
|---|---|---|---|---|
| | | | | BACHD vs. BACHD *Pias1* siRNA padj=0.0042 |
| | | | | BACHD vs. dN17-BACHD padj>0.9999 |

**Supplementary Table 3:** Mitochondria DEPs, downstream analysis and PIAS1 DEGs from previous study [65]. Refer to .XLS File

**Supplementary video 1:** Representative cryoET tomogram of a neurite in an HD patient iPSC-derived neuron (Q77) containing a prominent mitochondrion with aberrantly enlarged granules in the mitochondrial matrix composed of tightly packed, heterogeneous densities. Segmentation colors: red: microtubules, yellow: mitochondrial double-membranes, dark blue: granules, and cyan: cristae membranes.

**Supplementary video 2:** Representative cryoET tomogram of a neurite in an HD patient iPSC-derived neuron (Q66) displaying a double membrane-bound compartment with a large sheet aggregate. Segmentation colors: orange: sheet aggregate, cerulean blue: double membrane.
